# Mapping Alzheimer’s Molecular Pathologies in Large-Scale Connectomics Data: A Publicly Accessible Correlative Microscopy Resource

**DOI:** 10.1101/2023.10.24.563674

**Authors:** Xiaomeng Han, Peter H. Li, Shuohong Wang, Tim Blakely, Sneha Aggarwal, Bhavika Gopalani, Morgan Sanchez, Richard Schalek, Yaron Meirovitch, Zudi Lin, Daniel Berger, Yuelong Wu, Fatima Aly, Sylvie Bay, Benoît Delatour, Pierre Lafaye, Hanspeter Pfister, Donglai Wei, Viren Jain, Hidde Ploegh, Jeff Lichtman

## Abstract

Connectomics using volume-electron-microscopy enables mapping and analysis of neuronal networks, revealing insights into neural circuit function and dysfunction. In Alzheimer’s disease (AD), where amyloid-β (Aβ) and hyperphosphorylated-Tau (pTau) are implicated, connectomics offers an approach to unravel how these molecules contribute to circuit alterations by enabling the study of these molecules within the context of the complete local neuronal and glial milieu. We present a volumetric-correlated-light-and-electron microscopy (vCLEM) protocol using fluorescent nanobodies to localize Aβ and pTau within a large-scale connectomics dataset from the hippocampus of the 3xTg AD mouse model. A key outcome of this work is a publicly accessible vCLEM dataset, featuring fluorescent labeling of Aβ and pTau in the ultrastructural context with segmented neurons, glia, and synapses. This dataset provides a unique resource for exploring AD pathology in the context of connectomics and fosters collaborative opportunities in neurodegenerative disease research. As a proof-of-principle, we uncovered new localizations of Aβ and pTau, including pTau-positive spine-like protrusions at the axon initial segment and changes in the number and size of synapses near Aβ plaques. Our vCLEM approach facilitates the discovery of both molecular and structural alterations within large-scale EM data, advancing connectomics research in Alzheimer’s and other neurodegenerative diseases.

## Introduction

Alzheimer’s disease (AD) and related dementias are escalating global health crises, affecting over 6 million people in the U.S., with care costs projected to surpass $1 trillion annually by 2050 (“2024 Alzheimer’s Disease Facts and Figures” 2024). The annual incidence of dementia is expected to nearly double by 2060, underscoring the urgent need to uncover mechanisms underlying AD pathology and develop more effective treatments (M. Fang et al. 2025). Despite decades of research targeting pathological markers such as amyloid-β (Aβ) and hyperphosphorylated tau (pTau), most therapeutic efforts have achieved only modest efficacy (Cummings et al. 2024).

Traditional studies of AD often focus on isolated aspects—gene expression changes, protein alterations, or synaptic density—without capturing how these abnormalities interact within neuronal networks, a key driver of cognitive impairments (Yu, Sporns, and Saykin 2021). The field has been asking for integrative approaches that can access perturbation of the diseases at molecular, cellular as well as anatomical levels (Urbanski et al. 2019; Wood, Winslow, and Strasser 2015). Volume electron microscopy (volume EM) (Peddie et al. 2022)-based connectomics (Swanson and Lichtman 2016) offers a transformative framework for studying these interactions by providing nanoscale insights into neuronal and synaptic organization while simultaneously revealing cellular and subcellular pathology. This approach, while not yet extensively applied to AD, has been used in human tissue studies (Loomba et al. 2022; Shapson-Coe et al. 2024; Karlupia et al. 2023) and holds promise for advancing AD research.

Volume EM techniques, such as focused ion beam-scanning electron microscopy (FIB-SEM) (Knott et al. 2008), have provided valuable insights into synapse morphology and density in AD samples (Montero-Crespo et al. 2021; Domínguez-Álvaro et al. 2018; Blazquez-Llorca et al. 2013). However, their limited imaging volume (∼10 µm³) restricts the study of larger networks. Advances in ATUM-SEM (Hayworth et al. 2006; Kasthuri et al. 2015) and TEMCA (Lee et al. 2016) have enabled datasets on the millimeter scale, encompassing thousands of neurons and more comprehensive circuits (MICrONS Consortium et al. 2021; Loomba et al. 2022; Motta et al. 2019; Sievers et al. 2024; Nguyen et al. 2022; Kuan et al. 2024; Shapson-Coe et al. 2024). Yet, datasets specific to AD remain small and fragmented, hindering a holistic understanding of the disease. A hybrid method, ATUM-FIB, combines large-scale context information with local high-resolution FIB volumes and has been applied to samples from the AD mouse model 5xFAD (Kislinger, Fabig, et al. 2023). Despite these advancements, existing Alzheimer’s datasets remain small and fragmented, with the largest ATUM-SEM dataset for AD models covering only ∼45,000 cubic microns (Jiang et al. 2022; Pang et al. 2022; Kislinger, Niemann, et al. 2023). The scarcity of larger-scale vEM datasets restricts the ability to comprehensively analyze and interpret alterations in neuronal networks and cellular structures, which are critical for understanding the mechanisms of AD and related dementias.

In addition to neuronal connectivity disruptions, AD is characterized by pathological protein aggregates, including amyloid-β (Aβ) (Taylor, Hardy, and Fischbeck 2002; G.-F. Chen et al. 2017) and hyperphosphorylated tau (pTau) (Mandelkow and Mandelkow 1998; Noble et al. 2013) in the brain parenchyma. Despite their importance, there is very limited knowledge of how these hallmark pathological proteins exhibit distinct and identifiable ultrastructural features in EM datasets, making additional molecular labeling essential for their reliable detection. Extracellular Aβ accumulates into dense-core or diffuse plaques, which are detectable by EM (Blazquez-Llorca et al. 2013), while intracellular Aβ— potentially more neurotoxic than plaques—has also been identified (Gouras et al. 2010; LaFerla, Green, and Oddo 2007; Kelly et al. 2021). However, the precise manifestation of intracellular Aβ in EM images remains elusive, underscoring the importance of complementary molecular techniques. Similarly, pTau aggreagtes, identifiable by TEM as either straight or paired helical filaments (Crowther 1991), localize to cytoplasm, dendrites, and axons. Soluble forms of pTau are considered more neurotoxic than aggregates (Noble et al. 2013), yet whether either form exhibits specific ultrastructural characteristics detectable in ATUM-SEM data remains uncertain. These gaps highlight the critical need for innovative molecular labeling strategies to advance the study of AD pathology within large-scale EM datasets. Traditional methods for detecting these proteins often rely on detergents, which compromise sample ultrastructure (Im et al. 2019). Advanced labeling techniques are necessary to localize these pathological proteins reliably while preserving ultrastructure.

To address these challenges, we employed volumetric correlated light and electron microscopy (vCLEM), combining detergent-free immuno-probe (nanobody or single-chain variable fragment, scFv) - assisted immunofluorescence with large-scale ATUM-SEM imaging (T. Fang et al. 2018; Han et al. 2024). Nanobodies and scFvs, being only one-tenth or one-fifth the size of full-length IgG antibodies (see Figure 1 a for nanobody), exhibit excellent diffusion properties, enabling efficient labeling of intracellular proteins without requiring detergents. This approach preserves the ultrastructure of the sample (compare Figures 1 b and 1 c) and allows the labeling of multiple proteins in thick, unpermeabilized tissue. The labeled protein localizations can then be superimposed onto EM data generated by ATUM-SEM from the same sample, providing a powerful tool to directly link protein alterations to connectomic data. Despite its potential, vCLEM techniques have not yet been applied to AD or other dementia-related samples to facilitate research into these diseases.

**Figure 1.**
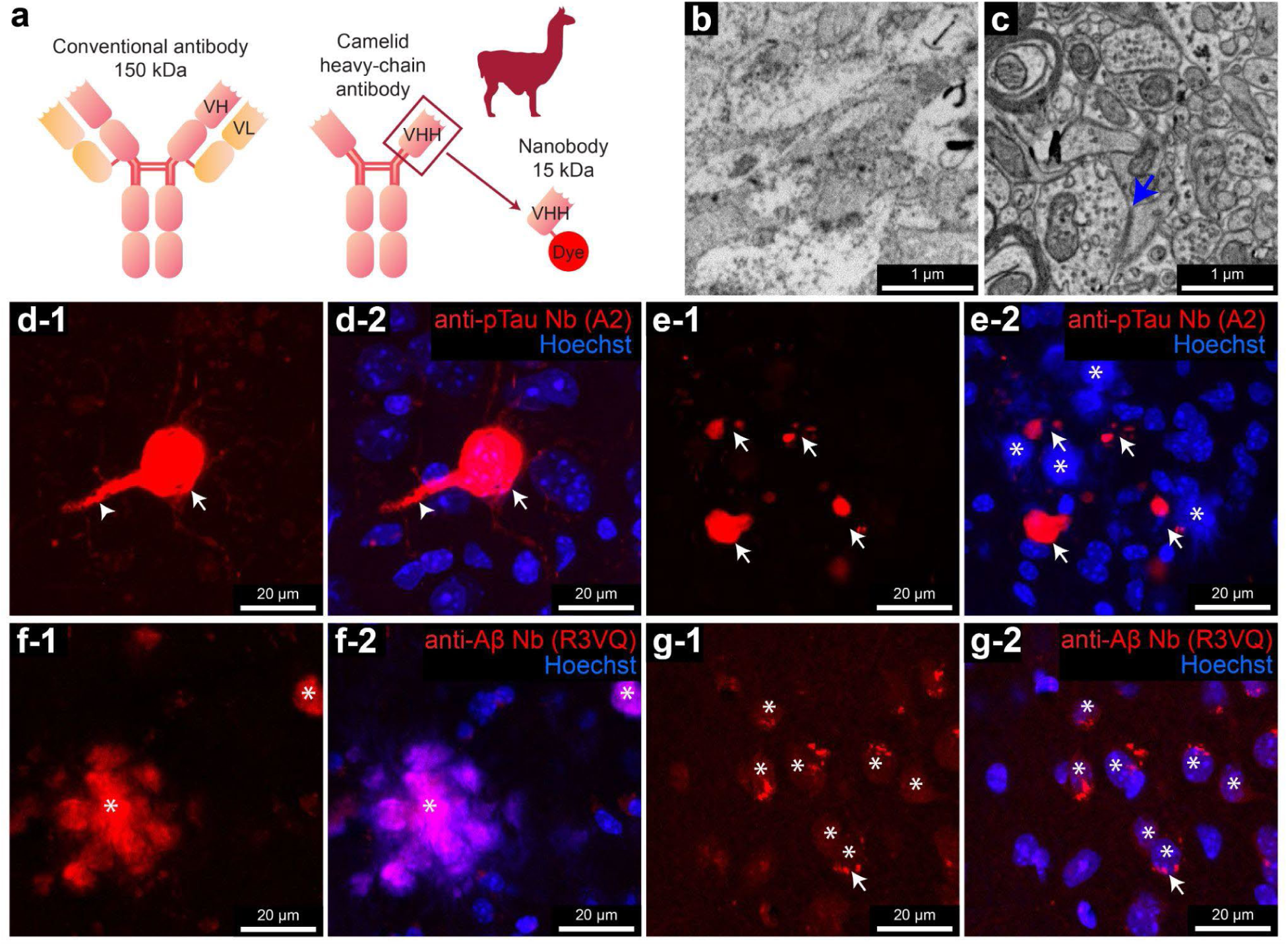
Characterization of fluorescent nanobody probes for pTau and Aβ. **a**, Schematic representations of a full-length IgG antibody, a camelid antibody, and a nanobody probe with a conjugated fluorescent dye. **b**-**c**, Comparison of ultrastructure between a sample (**b**) treated by Triton X-100 and a sample (**c**) labeled by nanobody probes, untreated with Triton X-100. Blue arrow in **c** indicates a synapse. **d-1** to **e-2**, Confocal images from the hippocampus of a 3xTg mouse labeled with a pTau-specific nanobody probe (A2) conjugated with the red dye 5-TAMRA. The panels **d-1**, **e-1** only show the nanobody labeling. Arrows in **d-1** and **d-2** show the labeled neuronal cell body. Arrowheads in **d-1** and **d-2** show the labeled apical dendrite. Arrows in **e-1** and **e-2** show the labeled neurites. Asterisks in **e-2** show extracellular plaque material labeled by Hoechst, as reported before (Uchida and Takahashi 2008). **f-1** to **g-2**, Confocal images from the hippocampus of a 3xTg mouse labeled with an Aβ-specific nanobody probe (R3VQ) conjugated with the red dye 5-TAMRA. The panels **f-1**, **g-1** only show the nanobody labeling. Asterisks in **f-1** and **f-2** show the labeled plaque material. Asterisks in **g-1** and **g-2** show the labeled neuronal cell bodies. Arrows in **g-1** and **g-2** show the autofluorescent lipofuscin.

In this study, we leveraged the vCLEM technique to localize Aβ and pTau in the hippocampal CA1 region of a female 3xTg AD model mouse. We developed two fluorescent nanobody probes targeting Aβ and pTau, which proved effective for detergent-free immunofluorescence. Alongside additional probes for microglia/macrophages (CD11b), astrocytes (GFAP), and blood vessels (Ly-6C/6G), these enabled precise molecular labeling of thick, unpermeabilized brain tissue. Using this approach, we generated and share <u>a large-volume ATUM-SEM-based connectomics dataset</u> with preserved ultrastructure, integrating molecular and structural information. In the dataset, we were able to localize extracellular Aβ plaques, intracellular Aβ and pTau at various novel locations.

Machine learning-based segmentation, including style transfer (Michał Januszewski et al. 2018; Michal Januszewski and Jain 2019) and supervised learning (Lin et al. 2021), facilitated automatic segmentation of neurons, dendrites, axons, glial processes, and synapses. The fully segmented dataset, now publicly accessible, supports 3D reconstruction of neurons, glia, their compartments, and synapses. Importantly, it is designed to be user-friendly for researchers in the AD field who may not be familiar with volume EM-based connectomics, enabling detailed analyses of cellular and network alterations. As a proof of principle, we identified previously unreported pTau localizations in enlarged, spine-like structures at the axon initial segment and observed synaptic changes surrounding an Aβ plaque. These findings highlight the potential of our dataset to uncover novel pathologies at molecular, cellular, and circuit levels.

## Results

### Generation of fluorescent nanobody probes for detergent-free immunolabeling of pTau and Aβ

To verify that detergent-free immunolabeling using nanobodies can localize pTau and Aβ in a vCLEM-compatible manner, we adapted two previously developed and validated nanobodies— A2, which specifically binds tau protein phosphorylated at multiple sites including S422; R3VQ, which selectively binds Aβ40/42 without cross-reacting with amyloid precursor protein (APP), the precursor from which Aβ is derived (T. Li et al. 2016; Lafaye et al. 2019, 2020)—into fluorescent probes (see Table 1). To facilitate broader use, we designed expression plasmids of these two nanobodies, optimized for easy, high-yield purification by Expi 293 cells and direct fluorescent dye conjugation (see Methods). These plasmids have been deposited at Addgene, and we provide a step-by-step protocol (Sup. File 1) to enable the generation of these probes.

**Table 1.** Fluorescent nanobody immuno-probes generated in this work.

| Target | Clone No. | Source | Fluorescent dyes conjugated | Link for expression plasmid at Addgene |
| --- | --- | --- | --- | --- |
| pTau | A2 | (Li et al. 2016) | 5-TAMRA, Alexa Fluor 594, Alexa Fluor 647 | https://www.addgene.org/233218/ |
| Aβ 40/42 | R3VQ | (Li et al. 2016) | 5-TAMRA, Alexa Fluor 647 | https://www.addgene.org/233217/ |

We tested two red fluorescent (5-TAMRA-conjugated) A2 and R3VQ probes using our detergent-free immunofluorescence protocol (see Methods) on tissue sections from the hippocampus CA1 region of 3xTg mice that preserves fine ultrastructure and is compatible with vCLEM (Compare Figure 1 b and c). This AD mouse model shows neurofibrillary tangles formed by pTau, Aβ extracellular plaques and intraneuronal Aβ deposits (Oddo et al. 2003).The A2 probe selectively labeled neurofibrillary tangle (NFT)-like structures in the cell bodies and dendrites of certain neurons (Figure 1 d-1, d-2; Sup. Fig. 1), as well as the dystrophic neurites nearby plaque material (Figure 1 e-1, e-2; Sup. Fig. 1). The R3VQ probe selectively labeled extracellular plaques (Figure 1 f-1, f-2; Sup. Fig. 1) and the cell bodies of certain neurons (Figure 1 g-1, g-2; Sup. Fig. 1).

**Figure 2.**
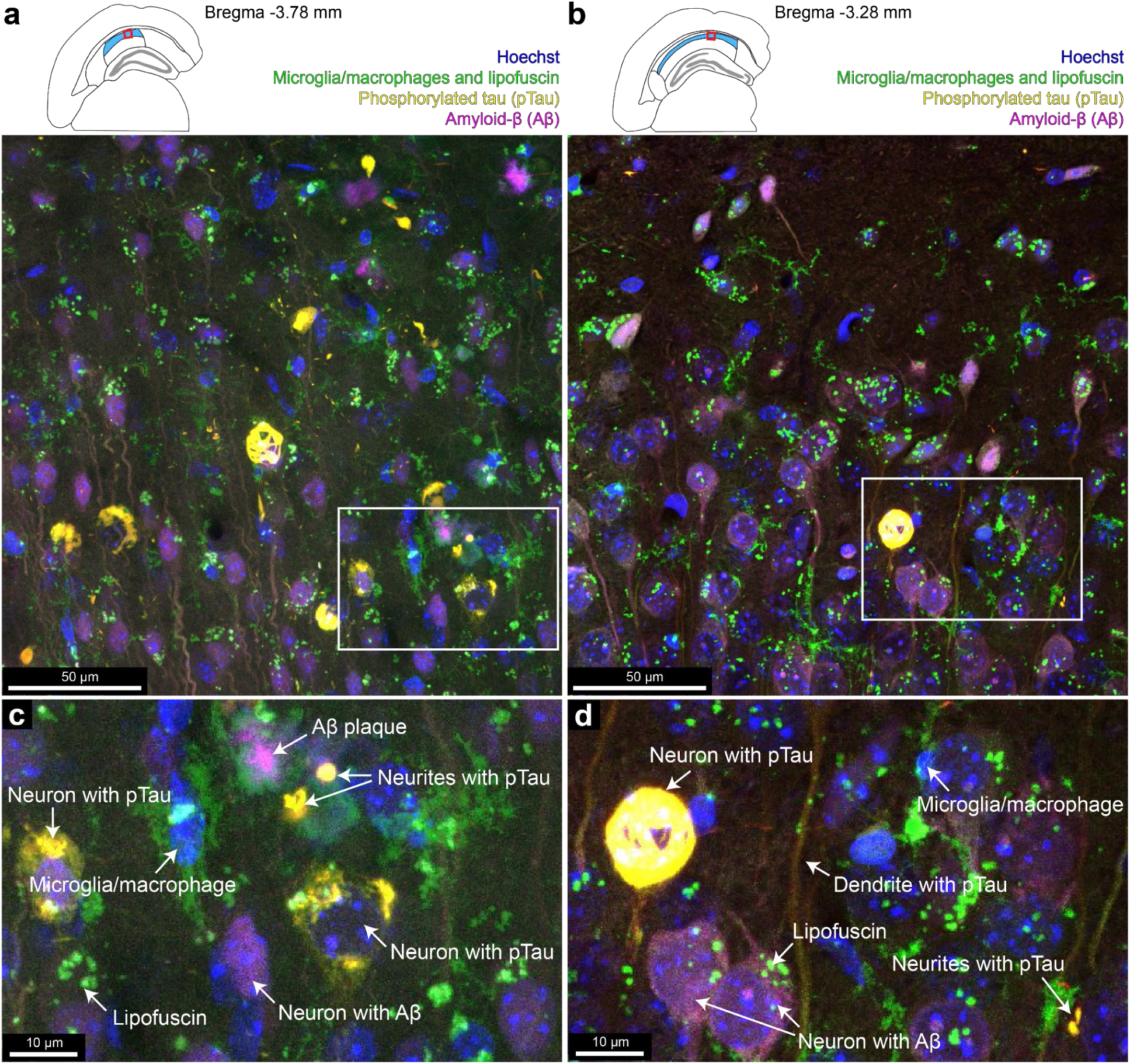
Four-color detergent-free immunofluorescence enabled by nanobody probes. Maximum intensity projection of the multi-color fluorescence image stacks acquired from the hippocampus at the Bregma −3.78 mm (**a**) or the Bregma −3.28 mm (**b**) of a 3xTg mouse labeled with a CD11b-specific nanobody probe conjugated with Alexa Fluor 488, a pTau-specific nanobody probe conjugated with 5-TAMRA, and an Aβ-specific nanobody probe conjugated with Alexa Fluor 647. Red boxes in the coronal section illustrations show where the image stacks were acquired. The stratum pyramidale of the hippocampal CA1 region is labeled in blue in the illustrations. **c**-**d**, enlarged boxed insets from **a**-**b**. Because the red fluorescence signal from the anti-pTau probe was far stronger than the green signal from the anti-CD11b probe, we saw crosstalk in the green channel. This meant that in the overlay, the pTau label appears yellow whereas the microglia appear green, but the two labels were nonetheless unambiguous. Another issue was that aged mice have lipofuscin particles in cell bodies of neurons and glia. This non-specific fluorescence localized to inclusion bodies that were easily differentiated from the membrane labeling of microglia.

Interestingly, we also observed R3VQ labeling in neuronal nuclei within neurons showing intracellular R3VQ signal (Figure 1 g-1, g-2; Sup. Fig. 1 e). While nuclear localization of Aβ is not widely established, a few reports suggest it may occur under certain conditions (Barucker et al. 2014; D’Andrea et al. 2021). However, given that prolonged fixation with glutaraldehyde was used in our protocol to preserve ultrastructural, this nuclear signal could also be an artifact of fixation-induced autofluorescence or redistribution of Aβ. Further investigation, such as alternative fixation protocol and colabeling approaches, would be required to clarify this observation.

Double immunofluorescence using A2 and anti-pTau mAb AT8, and R3VQ and anti-Aβ mAb 4G8 confirmed colocalization of signals (Sup. Figure a-1 to a-3, b-1 to b-3). No labeling was detected with A2 or R3VQ in sections from age-matched control animals (Sup. Figure 1 d and f). These results demonstrate that these fluorescent nanobody probes can localize Aβ and pTau in a detergent-free immunolabeling approach compatible with vCLEM.

Leveraging the flexibility of our protocol for easy conjugation of different fluorophores to the nanobodies, a broader application of these probes is for multiplex detergent-free immunolabeling. As a proof-of-principle, we combined these two probes with three additional nanobody probes for multiplex detergent-free immunolabeling. In one set of experiments, We used three fluorescent nanobody probes (A2, R3VQ, and anti-CD11b) to label two 120-μm thick brain sections in the absence of detergent (Figure 2 a, b; see Sup. Figure 2 for the anatomical details of the labeled sections) of a one-year-old female 3xTg mouse. CD11b is a marker for myeloid cells, including microglia and macrophages in the brain parenchyma (Bisht et al. 2016; Sedgwick et al. 1991; Martin et al. 2017). We acquired a four-color fluorescence 57-μm image stack (Figure 2 a) from the first section and a 21-μm stack from the second section (Figure 2 b; see Sup. Figure 2 for the anatomical details of the imaged regions of interest) in the stratum pyramidale in the CA1 region of the hippocampus. In both image stacks, NFT-like structures in the cell bodies and dendrites of certain pyramidal neurons and some dystrophic neurites were labeled by the anti-pTau probe (yellow signals in Figure 2 c, d). In addition, intraneuronal Aβ of certain pyramidal neurons were labeled by the anti-Aβ probe (magenta signals in Figure 2 c, d). Microglia/macrophages were labeled by the anti-CD11b probe (green signals in Figure 2 c, d). We noted that these labeled structures/cells were more abundant, and the labeled apical dendrites appeared to be more tortuous in the image stack acquired from the more caudal section (Figure 2 c, d). Only in this image stack (Figure 2 a, c) did we find five extracellular Aβ plaques labeled by the anti-Aβ 40/42 probe (magenta signals in Figure 2 a, c). These findings are consistent with the fact that the pathologies of neurofibrillary tangles and Aβ plaques progress from caudal to rostral in the hippocampus in the 3xTg mice (Belfiore et al. 2019).

In another experiment, we used four fluorescent nanobody probes (A2, anti-CD11b, anti-GFAP, and anti-Ly-6C/6G) to label a 120-μm thick brain section at Bregma −3.78 mm of another one-year-old female 3xTg mouse in the absence of detergent. Ly-6C is a lymphocyte antigen expressed by circulating monocytes recruited to sites of inflammation and endothelial cells of blood vessels (Martínez-Carmona et al. 2021; Mildner et al. 2007). We acquired a five-color fluorescence image stack of 23 μm (Figure 3).

**Figure 3.**
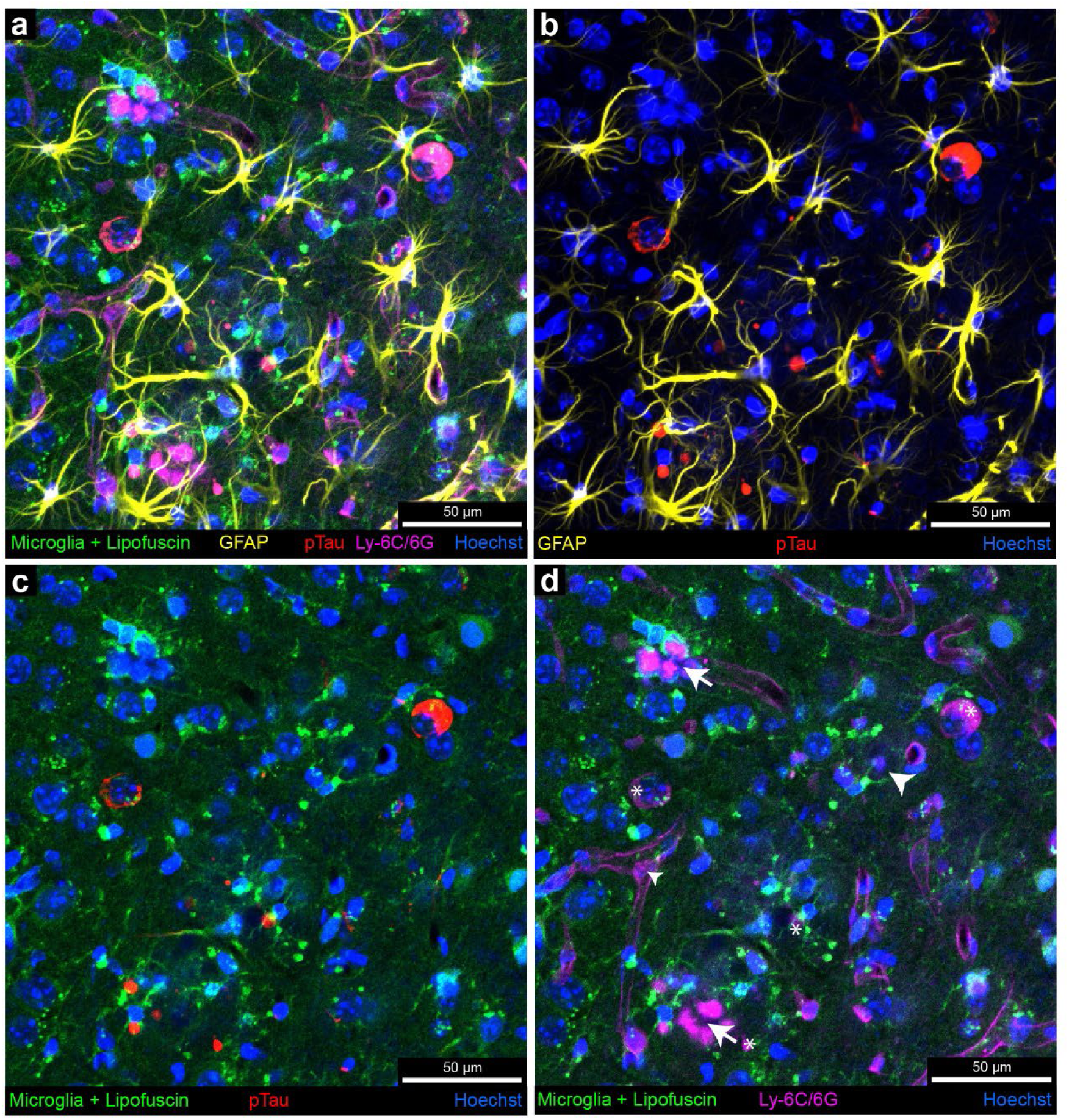
Five-color detergent-free immunofluorescence enabled by nanobody probes. **a**, Maximum intensity projection of the multi-color fluorescence image stacks acquired from the hippocampus of a 3xTg mouse labeled with a CD11b-specific nanobody probe conjugated with Alexa Fluor 488, a GFAP-specific nanobody probe conjugated with 5-TAMRA, a pTau-specific nanobody probe conjugated with Alexa Fluor 594, and a Ly-6C/6G-specific nanobody probe conjugated with Alexa Fluor 647. The signal of each fluorescent dye was pseudo-colored for better visualization. **b**-**d**, Three-channel maximum intensity projection images of the multi-color fluorescence image stack. White arrows in **d** show the labeled plaque material by the anti-Ly-6C/6G nanobody probe. The arrowhead shows plaque material not labeled by the anti-Ly-6C/6G nanobody probe. Asterisks show the crosstalk signals from the anti-pTau labeling.

Numerous activated astrocytes, labeled by the anti-GFAP nanobody probe, were observed nearby extracellular plaque material and neurons/neurites containing pTau (yellow signals in Figure 3 a, b). Interestingly, the anti-Ly-6C/6G nanobody labeled not only blood vessels but also some, though not all, plaque material (white arrows in Figure 3 d), which was surrounded by microglia/macrophages marked by the anti-CD11b probe (green signals in Figure 3 d). Since Ly-6C labels monocytes recruited to inflammation sites, which differentiate into macrophages also expressing CD11b (Mildner et al. 2007), it is likely that the CD11b-positive cells surrounding plaques are macrophages derived from monocytes, rather than resident microglia.

### Multi-color, large-scale vCLEM dataset with nanobody labeling, segmented neurons, glia and synapses

From the section shown in Figure 2 a, we acquired a high-resolution vEM dataset measuing 245 μm x 238 μm x 23 μm at a resolution of 4 nm x 4 nm x 30 nm (Figure 4 a) through ATUM-multi-SEM (see Methods). To the best of our knowledge, this represents the largest vEM volume obtained from an AD mouse model. Notably, the ultrastructure was well preserved across the entire volume, attributed to the exclusion of detergents during labeling, enabling the clear identification of various types of synapses (Figure 4 b, blue arrows).

**Figure 4.**
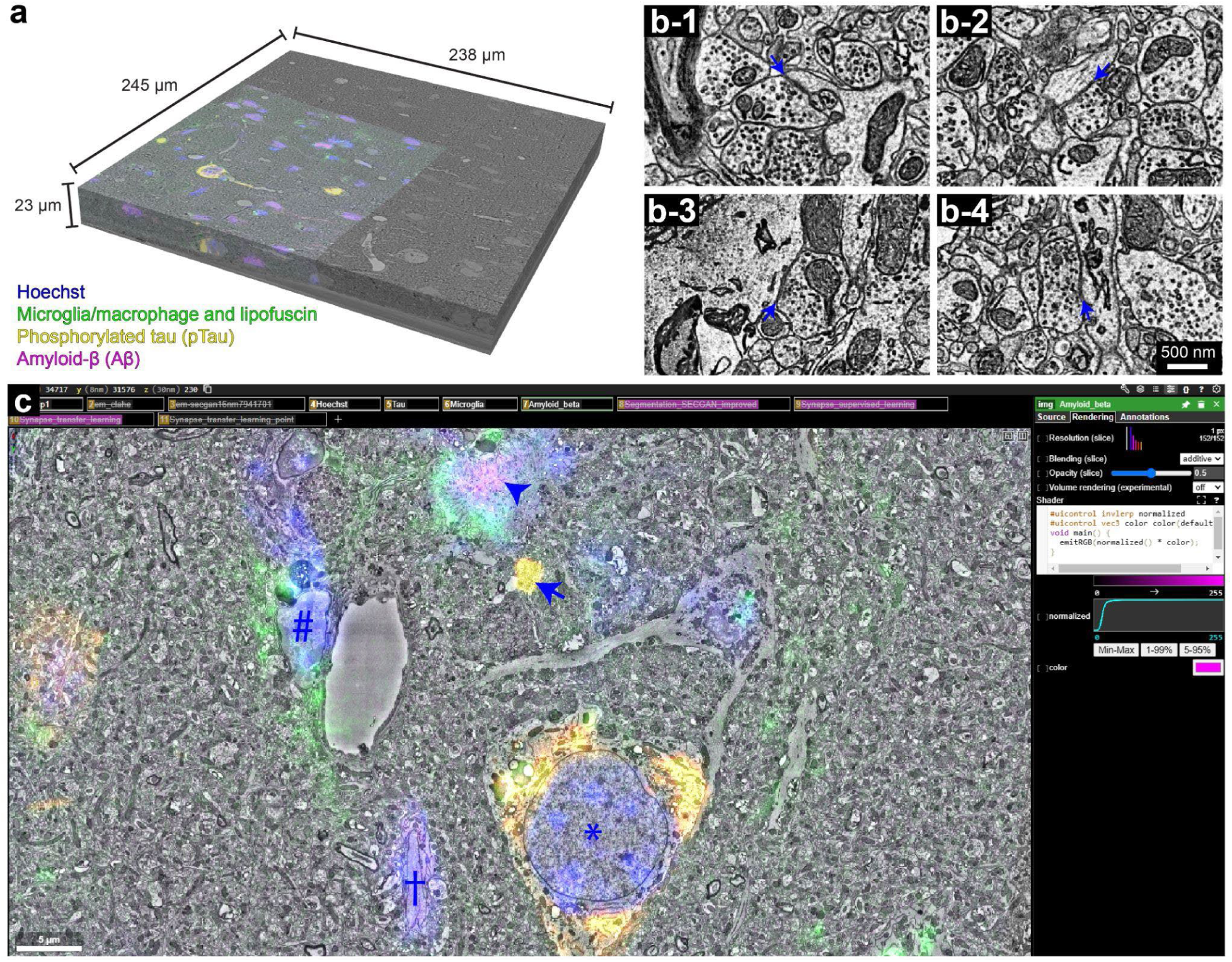
The four-color vCLEM dataset reveals ultrastructural abnormalities that colocalize with nanobody labeling. **a**, The high-resolution vEM volume acquired from the hippocampus CA1 region with four-color immunofluorescence from nanobody probes. The four-color fluorescence data was co-registered with the high-resolution vEM data. **b-1** to **b-4**, the preserved ultrastructure from this vEM volume. The blue arrows show synapses on a dendritic spine head (**b-1**), a dendritic shaft (**b-2**), a neuronal cell body (**b-3**), and an axon initial segment (**b-4**). **c**, Visualization of the vCLEM dataset in the Neuroglancer interface, overlaying fluorescence and EM ultrastructure. The asterisk indicates the cell body of a pyramidal neuron labeled by the anti-pTau nanobody. The arrow indicates the dystrophic neurite labeled by the anti-pTau nanobody. The arrowhead indicates the plaque material labeled by the anti-Aβ nanobody. The obelisk indicates the cell body of a pyramidal neuron labeled by the anti-Aβ nanobody. The hash mark indicates the microglia labeled by the anti-CD11b nanobody. This region corresponds to the one shown in Figure 2 c.

To enable the precise co-registration between the confocal fluorescence volume and the vEM dataset for generating a vCLEM dataset, the heavy-metal-stained, resin-embedded section was scanned by X-ray CT (microCT) (see Methods) prior to ATUM sectioning. The microCT image volume revealed regions with Aβ plaques and neuronal degeneration consistent with previous report (Barbone et al. 2022)(Sup. Figure 3). High-precision co-registration between the fluorescence volume and the vEM dataset was achieved by identifying corresponding landmarks, such as blood vessels and cell nuclei, across both volumes. We employed the FIJI/ImageJ plugin BigWarp (Bogovic et al. 2016) to perform a 3D transformation of the fluorescence volume via thin-plate spline interpolation to co-register the volumes. The transformed fluorescence volume, overlaid onto more than one-third of the high-resolution EM volume, is shown in Figure 4 a. This co-registration allows for the precise localization of fluorescence signals from each nanobody labeling within EM ultrastructure. In Figure 4 c, a representative region demonstrates that pTau (yellow) overlaps with the cell body of a pyramidal neuron and a dystrophic neurite, Aβ (magenta) associates with an Aβ plaque and the cell body of another pyramidal neuron, and CD11b (green) localized to a microglial cell/macrophage. We ingested this vCLEM dataset into Neuroglancer (Jeremy Maitin-Shepard, et al.), providing a publicly accessible <u>link</u> for users to visualize the fluorescence and the EM channels, either separately or in combination, and navigate through slices at different resolution levels (Figure 4 c; <u>Video 1</u>).

Efficient connectomic analysis of large vEM dataset critically depends on the rapid and accurate segmentation of image data. We employed flood-filling networks (FFNs) (Michał Januszewski et al. 2018) trained on a human cortex dataset (Shapson-Coe et al. 2024). While the direct application of the pre-trained FFNs was sufficient for segmenting neuronal cell bodies and dendrites, it performed suboptimally for segmenting axonal and glial processes (Sup. Figure 4 a). To address this, we utilized Segmentation Enhanced CycleGANs (Michal Januszewski and Jain 2019) to adjust the visual appearance of this data to more closely match the H01 dataset prior to segmentation. This approach substantially improved the segmentation of axonal and glial processes (Sup. Figure 4 b; Figure 5 a). The updated segmentation results are also publicly available as a segmentation layer (“Segmentation_SECGAN_improved”) through the previously mentioned link. Users can perform 3D reconstructions of neuronal or glial cells by manually agglomerating segments, simply by selecting the automatically generated segments associated with the target cell (Figure 5 c; <u>Video1</u>).

**Figure 5.**
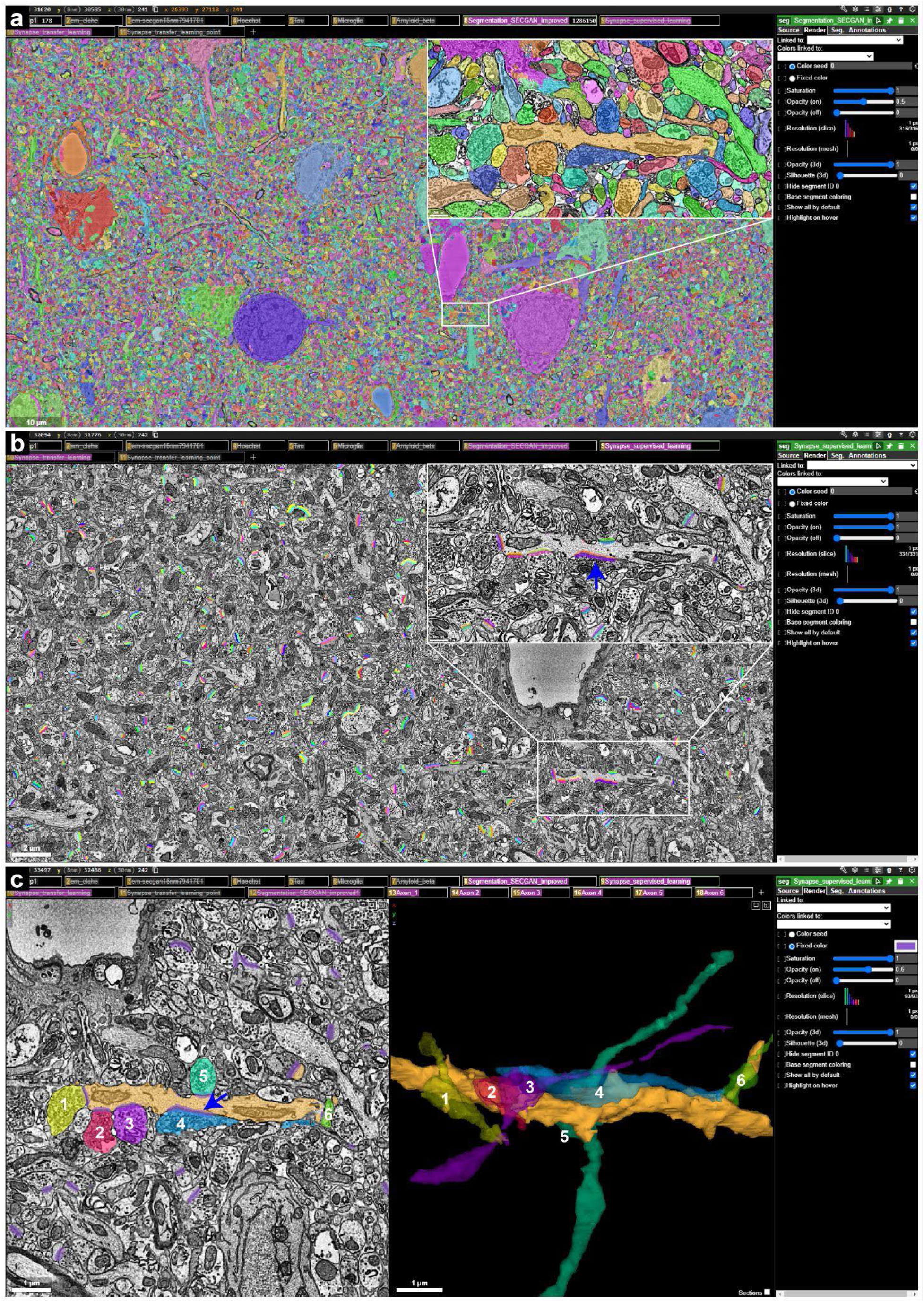
Efficient 3D reconstruction and synapse identification in the vCLEM dataset using segmentation and synapse prediction layers. **a**, Visualization of the 3D segmentation layer (“Segmentation_SECGAN_improved”) in the Neuroglancer interface. Each segmented object is uniquely color-coded, and users can click on these objects to reconstruct them in 3D. **b**,Synapse prediction layer (“Synapse_supervised_learning”) visualized in Neuroglancer. Synapses are labeled with distinct colors for pre-synaptic (e.g., purple) and post-synaptic (e.g., orange) appositions. The blue arrow highlights a synapse located on a dendritic shaft. Users can click on these appositions to view their 3D structure. **c**, Example 3D reconstruction of a dendrite (orange) and six axonal profiles (numbered 1–6, each with a distinct color) using the segmentation and synapse prediction layers. Synapses formed by these axons are shown in purple. The blue arrow points to synapses made by axon No. 4.

Another key requirement for connectomic analysis is the comprehensive annotation and segmentation of all synapses. This is particularly crucial in AD studies, given the well-documented synaptic alterations (Scheff and Price 2006; Montero-Crespo et al. 2021, 2020) and the observed correlation between cognitive decline and reduced synapse density (DeKosky and Scheff 1990; Davies et al. 1987; Tzioras et al. 2023). Our vCLEM dataset offers the necessary resolution to precisely locate and analyze synapses at the ultrastructural level (Figure 4 b-1 to b-4). While manual annotation for synapse analysis is straightforward, it is not feasible for large-scale studies. Machine-learning-based automated synapse detection (Çiçek et al. 2016; Parag et al. 2019; Lin et al. 2021; Buhmann et al. 2021) has been successfully applied to large-scale connectomics datasets (Shapson-Coe et al. 2024; Turner et al. 2022; Scheffer et al. 2020), demonstrating efficiency, objectivity, and accuracy.

For automatic synapse prediction, we employed the U-Net deep learning model (Parag et al. 2019) to identify the pre- and postsynaptic components of each synapse. Two approaches were explored: a supervised learning approach and a transfer learning approach, utilizing two existing synapse prediction frameworks (Lin et al. 2021; Shapson-Coe et al. 2024). For evaluation, we manually annotated all synapses within a small subvolume measuring 6 μm × 6 μm × 1.5 μm.

In the supervised learning approach, synapses were annotated within a randomly selected subvolume measuring 6 μm × 6 μm × 6 μm, distinct from the evaluation subvolume. This training dataset primarily consisted of neuropil and excluded regions containing blood vessels or cell bodies. Following the guidelines of an online tutorial (Lin et al. 2021), we trained the synapse prediction model and applied the watershed transform to convert the predictions into synapse instances (see Methods). On the evaluation subvolume, this model achieved a precision of 86.06% and a recall of 98.88%. Applied to the entire dataset, the model identified approximately 3.5 million synapses. This first version of synapse predictions is publicly available via the aforementioned repository as an independent data layer (“Synapse_supervised_learning”) (Figure 5 b). In this layer, each synapse is labeled with one presynaptic segment and one postsynaptic segment, assigned consecutive segment IDs (Figure 5 b). Users can identify synapses in EM images and visualize them in 3D by interacting with the corresponding segments (Figure 5 c; <u>Video 1</u>). Additionally, these 3D visualizations can be integrated with cell reconstructions for advanced analysis (Figure 5 c; <u>Video 1</u>).

While effective, the supervised approach missed many synapses due to the limited diversity in the small training dataset. To address this limitation, we adopted a transfer learning approach using a pretrained synapse prediction model developed for a human cortex dataset (Shapson-Coe et al. 2024). To adapt it to this dataset, we first ran the synapse models trained on the human cortex dataset on this data with SECGAN style-transfer applied to mimic the human dataset. This produced 0.225 million synaptic masks with low false positive rates, but with a significant number of false negatives (estimated at about 50%), particularly within neuronal processes (Sup Figure 5 a-1 to a-3) which require filtering. We then used these masks as model-produced ground truth to retrain the synapse model from scratch on this dataset without SECGAN. The retrained model produced synapses with significantly improved precision of 46% and recall of 52%. Applied to the entire dataset, the model’s predictions were filtered to retain only synapses associated with neuronal membranes, resulting in the identification of approximately 0.7 million synapses. This second version of synapse predictions is also publicly available as a separate data layer (“Synapse_transfer_learning”) (Sup Figure 5 b-2). Like the first version, it enables users to identify, visualize, and reconstruct synapses in 3D. This version proved particularly effective in detecting synapses missed by the supervised approach (Sup Figure 5 b-2), leveraging the pretrained model’s knowledge of synapse morphology in the human cortex. Since both synapse detection layers are accessible in Neuroglancer, users can combine them, resulting in a union that achieves greater precision than either model alone. This dual-layer approach serves as a complementary resource for synapse identification and analysis, ensuring a more comprehensive and accurate mapping of synaptic structures.

The publicly accessible dataset is among the largest vEM datasets generated for an AD model, offering unique opportunities for the scientific community. With fully segmented neuronal and glial cells, as well as synapses, it provides an invaluable resource for researchers, particularly those studying AD and other dementias who may be less familiar with volume EM-based connectomics. This innovative dataset enables the following: (1) pinpointing cells and cellular structures where Aβ and pTau are localized using overlapping fluorescence signal layers; (2) reconstructing affected neurons and glial cells at nanometer resolution with the 3D segmentation layer to understand how AD-related abnormalities manifest in cellular morphology; (3) exploring how the localization of Aβ and pTau impacts synaptic connectivity and neuronal networks using the synapse segmentation layer; (4) generating new hypotheses based on findings from the dataset to be tested with other methodologies; and (5) comparing findings with those reported in the literature or observed in other animal models or human tissues. By lowering the barrier to entry for researchers unfamiliar with volume EM or segmentation tools, this dataset fosters broader participation in connectomics-driven AD research and encourages cross-disciplinary collaboration.

### vCLEM dataset uncovers ultrastructural pathologies that colocalize with nanobody labeling

To demonstrate the potential of this dataset for advancing AD research, we conducted an analysis of sites labeled with each of the three molecular markers (pTau, Aβ, CD11b). This analysis revealed an array of previously uncharacterized ultrastructural abnormalities colocalize with each marker, providing insights into molecular and structural pathology at the nanoscale. These findings, along with their spatial coordinates in the dataset, are compiled in Table 2 and discussed in detail in the following paragraphs, providing a valuable resource for further investigation. Researchers can leverage this dataset for in-depth 3D reconstruction of affected cells and subcellular structures, enabling new avenues for identifying and understanding AD-related pathology.

**Table 2.** Ultrastructural Pathologies that Colocalize with Nanobody Labeling.

| Label | Location Type | Subtype | Other Ultrastructural Property | Coordinates |
| --- | --- | --- | --- | --- |
| pTau |  |  |  |  |
|  | Neuronal soma |  |  |  |
|  |  | Only pTau |  |  |
|  |  |  | pTau with straight filament | (26467, 24370, 155) |
|  |  |  | pTau with no straight filament, but short-filament | (28449, 33128, 151),<br>(35520, 32879, 177),<br>(19278, 27572, 51),<br>(20141, 28463, 466) |
|  |  | Both pTau and A-beta |  | (30106, 31592, 365),<br>(18693, 21208, 510) |
|  | Dendrite |  |  |  |
|  |  | Straight |  | (25988, 27500, 121),<br>(28382, 36337, 121),<br>(35212, 35235, 142) |
|  |  | Tortuous |  | (21209, 33200, 220),<br>(25698, 33034, 266),<br>(21539, 35309, 235) |
|  | Axon |  |  |  |
|  |  | Bleb-like structure from AIS |  | (35464, 30205, 439),<br>(34864, 30905, 460) |
|  |  | Bleb-like structure from myelinated axon |  | (28612, 18697, 157) |
|  |  | Swelling |  | (28301, 30383, 77) |
|  | Glia |  |  | (32846, 20364, 661) |
| **Aβ** | Plaque |  |  | (34331, 29527, 215),<br>(32240, 12443, 266),<br>(36268, 27060, 438),<br>(32071, 16340, 110),<br>(39797, 13403, 748) |
|  | Neuronal soma | Only Aβ |  | (23970, 30423, 166),<br>(33496, 33578, 365) |
| **CD11b** | Cell associated with A-beta plaques |  |  | (32878, 30971, 183),<br>(34740, 27790, 255),<br>(33988, 28188, 7),<br>(37528, 35367, 55),<br>(36983, 26638, 404) |
|  | Cell associated with dystrophic neurites |  |  | (24013, 20988, 186),<br>(31741, 19684, 293),<br>(37100, 25861, 659),<br>(29528, 12771, 599) |
|  | Cell adjacent to neuron with intraneuronal A-beta |  |  | (23113, 30303, 100) |

\*The coordinates can be entered into Neuroglancer as in Sup. Figure 6 to move to the location.

pTau: Our analysis revealed a diverse array of pTau labeling patterns across different ultrastructural contexts, highlighting a complex landscape of pTau-associated abnormalities. We found NFT-like structures from the labeling of the anti-pTau nanobody localized within cell bodies of seven pyramidal neurons and one glial cell (red fluorescence, see Figure 6 a-1 to d-2 for four examples for neurons, and j-1 and j-2 for the glial cell) among the ∼100 neuronal and glial cells in the volume where the fluorescence signals were co-registered. Only one of the seven pTau positive pyramidal cells (which did not show intraneuronal Aβ), showed pathological straight filament-like structures (Figure 6 a-1, a-2; Sup. Figure 7 a), which is one of the ultrastructural manifestations of neurofibrillary tangles (Crowther 1991). The other pTau positive pyramidal cells showed short filament-like structures in their cell bodies (Figure 6 b-2, c-2). We suspect that these are not pathological because in other pyramidal neurons free of pTau labeling, similar structures were seen (Sup. Figure 7 c-1, c-2). Two of the pyramidal cells also showed intraneuronal Aβ (magenta fluorescence, see Figure 6 d-1, d-2 for one example). These two neurons had the same morphological features as neurons that only had intraneuronal Aβ (see below). We also found NFT-like structures localized in dendrites that were either straight or tortuous (Figure 6 e-1 to f-2). Again, we did not find specific ultrastructural manifestations of the neurofibrillary tangle structures in these dendrites despite them being pTau positive (Sup. Figure 7 d-1, d-2). We examined four axonal sites with pTau labeling. Two of these sites consisted of pTau positive, bleb-like structures each filled with straight filaments and stemmed from the axon initial segment of a pTau positive pyramidal neuron (see Figure 6 g-1, g-2; see Sup. Figure 7 b for one of them). Another site showed an enlarged pTau positive swelling filled with lipofuscin granules near the axon initial segment of a different pTau positive pyramidal neuron (Figure 6 h-1, h-2). The fourth side contained a large pTau positive bleb-like structure filled with degenerating organelles, emerging from a myelinated axon (Figure 6 i-1, i-2), similar to what has been seen previously (Gowrishankar et al. 2015). The labeled glial cell had glial inclusions known as “coiled bodies” (Ferrer et al. 2024; T. Li et al. 2016) which were specifically labeled by the anti-pTau probe (Figure 6 j-1, j-2). Based on our observation that not all pTau-labeled sites exhibited characteristic ultrastructural features of neurofibrillary tangles (NFTs), we propose that certain NFT-like structures or pTau aggregates may lack visible ultrastructural representations. This underscores the importance of using correlative microscopy and labeling technique like ours to identify neurons harboring these aggregates in EM micrographs, as they may otherwise go undetected.

**Figure 6.**
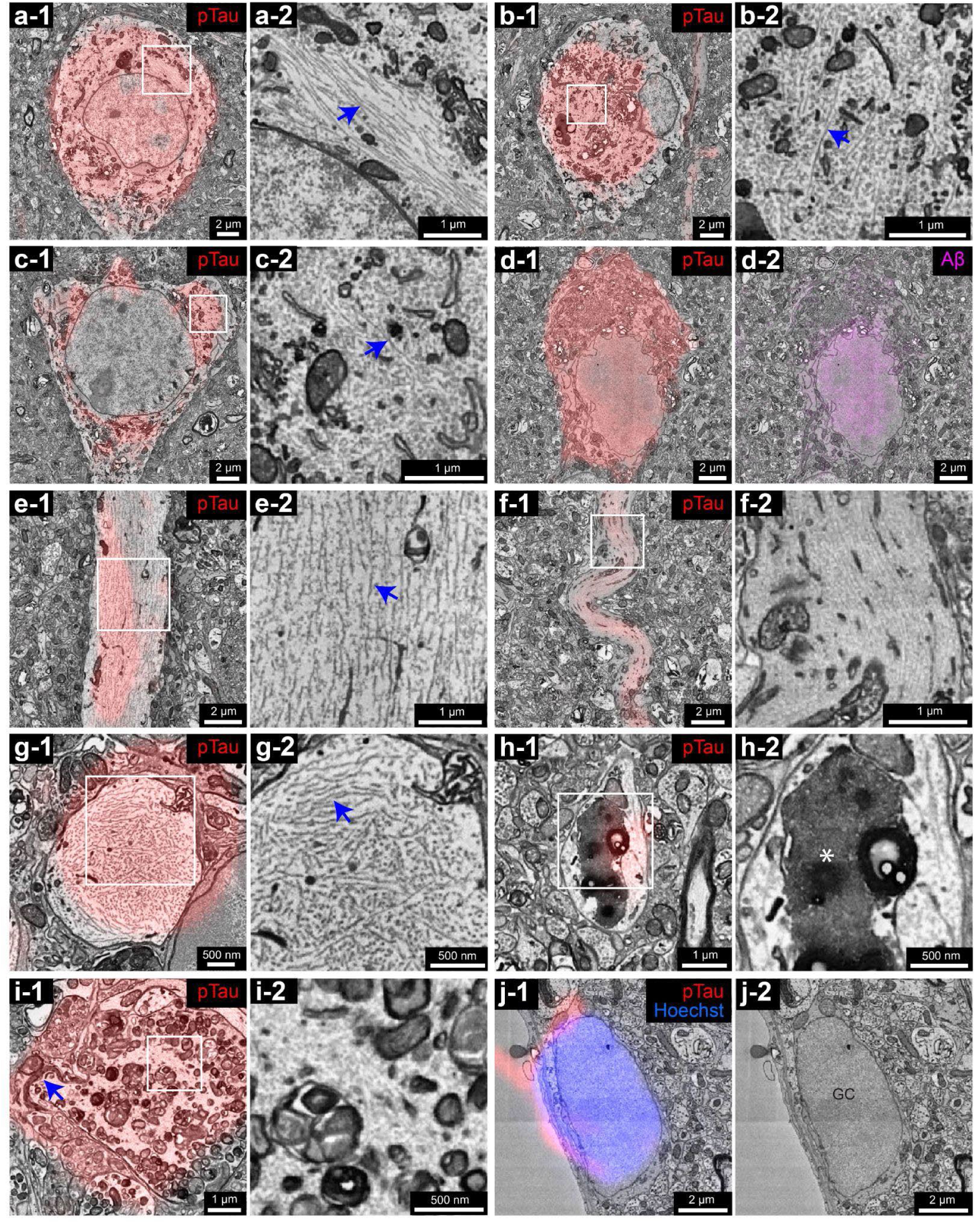
Ultrastructural abnormalities that colocalize with the labeling of pTau. **a-1**, red fluorescence from the anti-pTau nanobody overlaps with the cell body of a pyramidal neuron. **a-2**, enlarged inset from **a-1**. The blue arrow indicates the straight filament-like structure. **b-1** and **c-1**, red fluorescence from the anti-pTau nanobody overlaps with the cell bodies of two pyramidal neurons. **b-2** and **c-2**, enlarged insets from **b-1** and **c-1**. The red arrows indicate the short filament-like structures that were also seen in neurons that were not labeled by the anti-pTau nanobody (see Sup. Figure 4 c-1, c-2). **d-1** and **d-2**, the pyramidal neuron that was labeled by both the anti-pTau nanobody (red fluorescence in **d-1**) and the anti-Aβ nanobody (magenta fluorescence in **d-2**). **e-1** and **f-1**, red fluorescence from the anti-pTau nanobody overlaps with a straight apical dendrite and a tortuous apical dendrite of two pyramidal neurons. **e-2** and **f-2**, enlarged insets from **e-1** and **f-1**. The blue arrow in **e-2** indicates the short filament-like structure that was also seen in the apical dendrite that was not labeled by the anti-pTau nanobody (see Sup. Figure 4 d-1, d-2). **g-1**, red fluorescence from the anti-pTau nanobody overlaps with a bleb-structure connected to an axon. **g-2**, enlarged inset from **g-1**. The blue arrow indicates the straight filament-like structure. **h-1**, red fluorescence from the anti-pTau nanobody overlaps with an enlarged swelling of an axon filled with lipofuscin granules. **h-2**, enlarged inset from **h-1**. The asterisk shows the lipofuscin granules. **i-1**, red fluorescence from the anti-pTau nanobody overlaps with a bleb-like structure that emerged from a myelinated axon. This bleb was filled with degenerating organelles. The blue arrow indicates the end of the myelination. **i-2**, enlarged inset from showing the degenerating organelles. **j-1** and **j-2**, red fluorescence from the anti-pTau nanobody overlaps with a glial cell that had coiled bodies (red fluorescence in **j-1**).

Aβ: We found extracellular Aβ plaque structures from the labeling of the anti-Aβ nanobody (magenta fluorescence in Figure 7) localized within five separate spots of plaque material (see Figure 7 a-1 to b-2 for two examples) which showed the typical ultrastructure of dense-cored plaques (Blazquez-Llorca et al., 2013). These plaques had dense cores of Aβ deposits in the center, Aβ fibrils, glial processes and dystrophic neurites surrounding them. Intraneuronal Aβ localized within neurons that had very tortuous apical dendrites, shrunken cell bodies, dark osmiophilic cytoplasm and degenerating organelles (Figure 7 c-1 to d-2). These neurons with intraneuronal Aβ appeared to be more abnormal than neurons that only had pTau.

**Figure 7.**
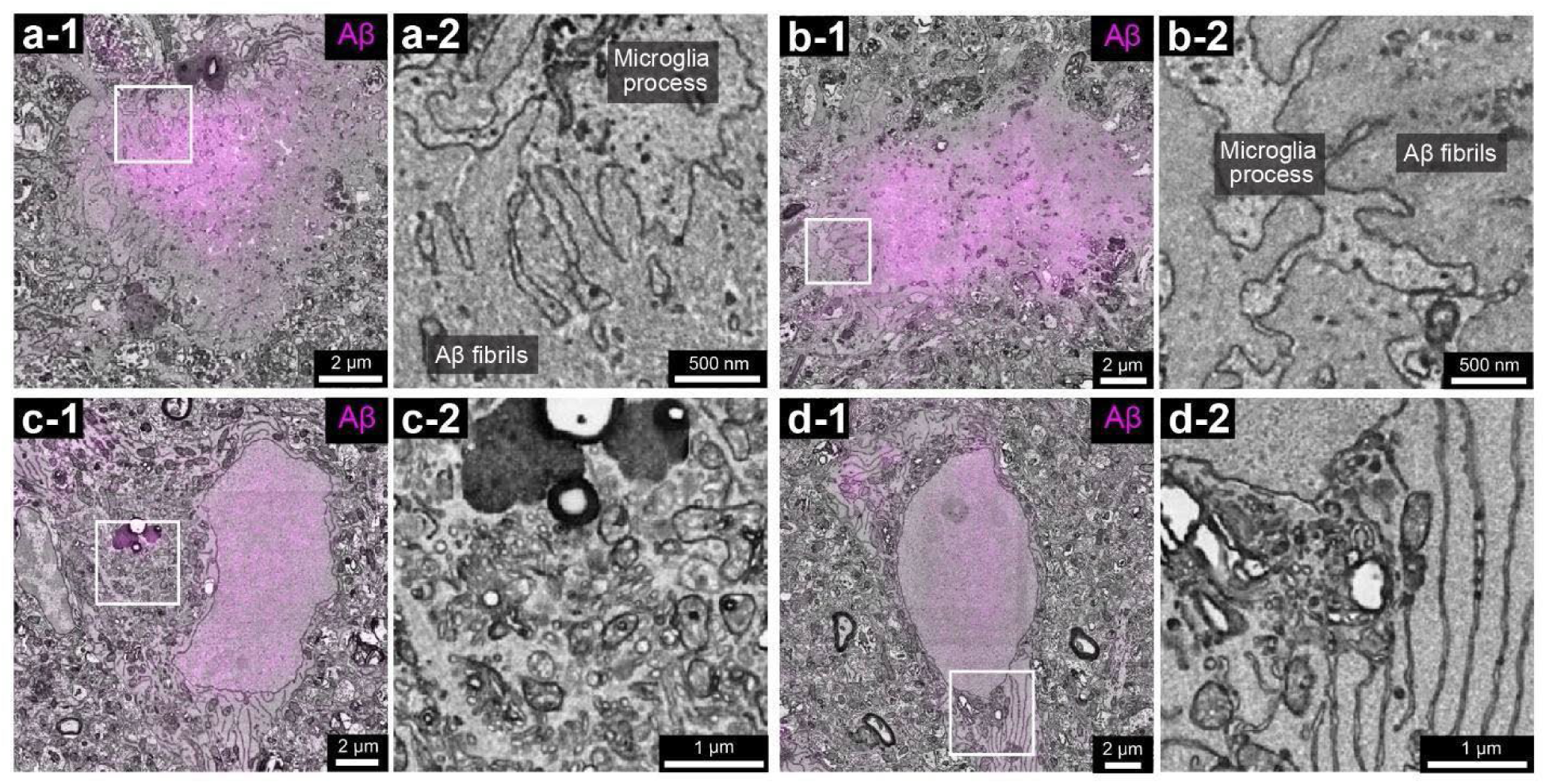
Ultrastructural abnormalities that colocalize with the labeling of Aβ. **a-1**, magenta fluorescence from the anti-Aβ nanobody overlaps with plaque material. **a-2**, enlarged inset from **a-1**. **b-1**, magenta fluorescence from the anti-Aβ nanobody overlaps with another plaque material. **b-2**, enlarged inset from **b-1**. **c-1**, magenta fluorescence from the anti-Aβ nanobody overlaps with the cell body of a pyramidal neuron. **c-2**, enlarged inset from **c-1** showing the degenerating organelles in this neuron. **d-1**, magenta fluorescence from the anti-Aβ nanobody overlaps with the cell body of another pyramidal neuron. **d-2**, enlarged inset from **d-1** showing the dilated endoplasmic reticula in this neuron.

CD11b: CD11b is a myeloid cell marker that labels both resident microglia and infiltrating macrophages within the brain parenchyma (Sedgwick et al. 1991; Martin et al. 2017). As a component of the phagocytic receptor CR3, CD11b is present on the plasma membrane of glial processes that surround synaptic elements (Bisht et al. 2016). Among these cells, some microglia, characterized by a condensed, electron-dense cytoplasm and nucleoplasm, were referred to as ‘dark microglia’ in a previous electron microscopy studies (Bisht et al. 2016; St-Pierre et al. 2022). In our vCLEM dataset, we identified ten labeled cells based on anti-CD11b nanobody labeling (green fluorescence in Figure 8; five examples shown), although we could not distinguish between microglia and macrophages. Among them, five were associated with Aβ plaques (Figure 8 a-1, c-3, c-4), extending ramified processes to encircle and perhaps phagocytose plaque material (Figure 8 b); four however were associated with dystrophic neurites where there was no Aβ plaque (Figure 8 c-1); one was adjacent to a neuron that had intraneuronal Aβ (Figure 8 c-2). These cells appeared stressed: they had a large amount of lipofuscin granules, osmiophilic cytoplasm, altered mitochondria, condensed nucleoplasm, and dilated endoplasmic reticulum (Figure 8 a-2, a-3), which may be attributed to the activation of the integrated stress response (ISR) as previously described (Flury et al. 2024).

**Figure 8.**
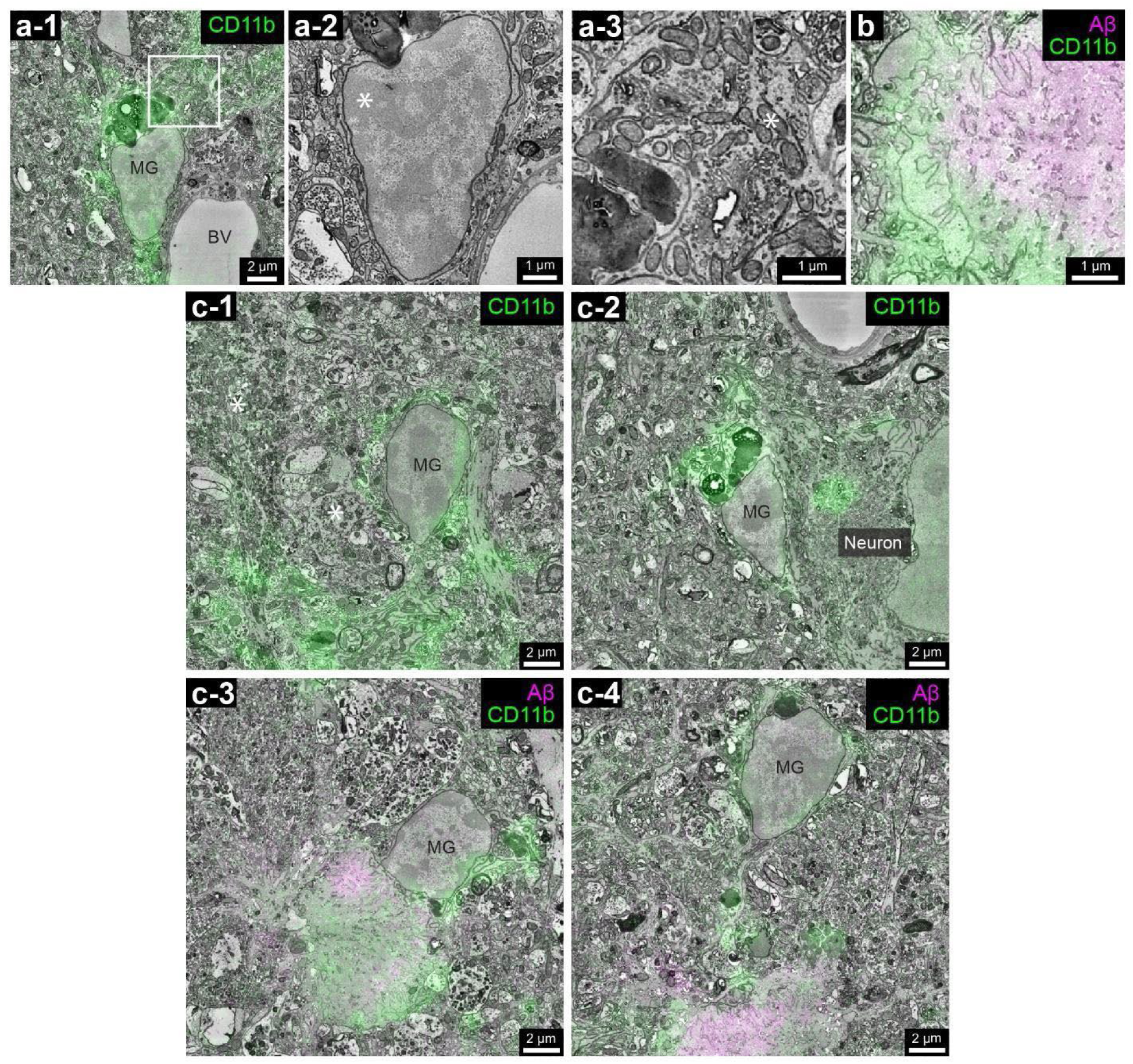
Ultrastructural abnormalities that colocalize with the labeling of CD11b. **a-1**, green fluorescence from the anti-CD11b nanobody overlaps with a microglial cell. **a-2**, the nucleus of the microglial cell in **a-1**. Asterisk indicates the condensed nucleoplasm. **a-3**, enlarged inset from **a-1**. Asterisk indicates the altered mitochondria. **b**, green fluorescence from the anti-CD11b nanobody overlaps with the glial processes encircling plaque material, which shows overlapped magenta fluorescence from the anti-Aβ nanobody. **c-1**, green fluorescence from the anti-CD11b nanobody overlaps with a microglial cell nearby dystrophic neurites (asterisks). **c-2**, green fluorescence from the anti-CD11b nanobody overlaps with a microglial cell next to a pyramidal neuron (the same neuron in Figure 6 c-1). **c-3** and **c-4**, green fluorescence from the anti-CD11b nanobody overlaps with two microglial cells nearby plaque material, which shows overlapped magenta fluorescence from the anti-Aβ nanobody.

### 3D Reconstructions Reveal Detailed Morphological Features of Neurons with Intraneuronal Aβ or pTau

As a proof-of-principle demonstration of the dataset’s potential, we conducted 3D reconstructions of neurons with intraneuronal Aβ or pTau, showcasing the dataset’s ability to uncover detailed, unreported morphological features associated with these pathologies. Using the automatic segmentation followed by manual proofreading, a pyramidal neuron with intraneuronal Aβ (Figure 7 c-1) and two pyramidal neurons with intraneuronal pTau (Figure 6 b-1, c-1) were reconstructed.

The cell body of neuron 1 with intraneuronal Aβ labeling appeared abnormal as if it were in the process of degenerating (Figure 7 c-1, c-2). A microglial cell was immediately adjacent raising the possibility that the neuron would soon be phagocytosed (Figure 8 c-2). Surprisingly, this neuron still retained multiple spiny dendrites (Figure 9 a, inset 1) and its axon projected normally for ∼100 μm without myelin and then showed normal myelination (Figure 9 a). Neuron 2 showed intraneuronal pTau labeling in its soma (Figure 6 b-1, b-2). Nonetheless it also had multiple normal looking spiny dendrites (Figure 9 a, inset 2). This neuron’s axon ran normally ∼80 μm before becoming myelinated (Figure 9 a). As was the case for the other two neurons, there were several spine-like structures innervated by vesicle-filled axonal profiles (Figure 9 b) along this neuron’s axon initial segment (AIS). This AIS also had an abnormally enlarged swelling that was positive for anti-pTau nanobody labeling. This swelling was filled with lipofuscin granules (see Figure 6 h-1, h-2). The third neuron (Neuron 3) with intraneuronal pTau labeling (Figure 6 c-1, c-2) was located such that its soma was adjacent to an Aβ plaque (Figure 9 a). This neuron also possessed multiple normal looking spiny dendrites (Figure 9 a inset 3) and an axon with normal appearance that ran ∼60 μm before becoming myelinated (Figure 9 a). Two spine-like swollen protrusions emerged from its AIS via narrow stalks (Figure 9 a box c, d-1, d-2; Figure 9 c). Each of these was filled with straight filaments, similar to those described above (Figure 9 d-1, d-2; also see Figure 6 g-1, g-2 and Sup. Figure 7 b). We suspect that these objects are abnormally enlarged postsynaptic spine heads because they were innervated by vesicle-filled axonal profiles (Figure 9 e-1, e-2). The protrusion stalks are extremely thin (less than 1 µm in diameter), suggesting the possibility that these structures may eventually detach from the AIS. Supporting this hypothesis, we identified a nearby pTau-filled bleb that appeared disconnected from the AIS or any adjacent structures (Sup. Figure 8).

**Figure 9.**
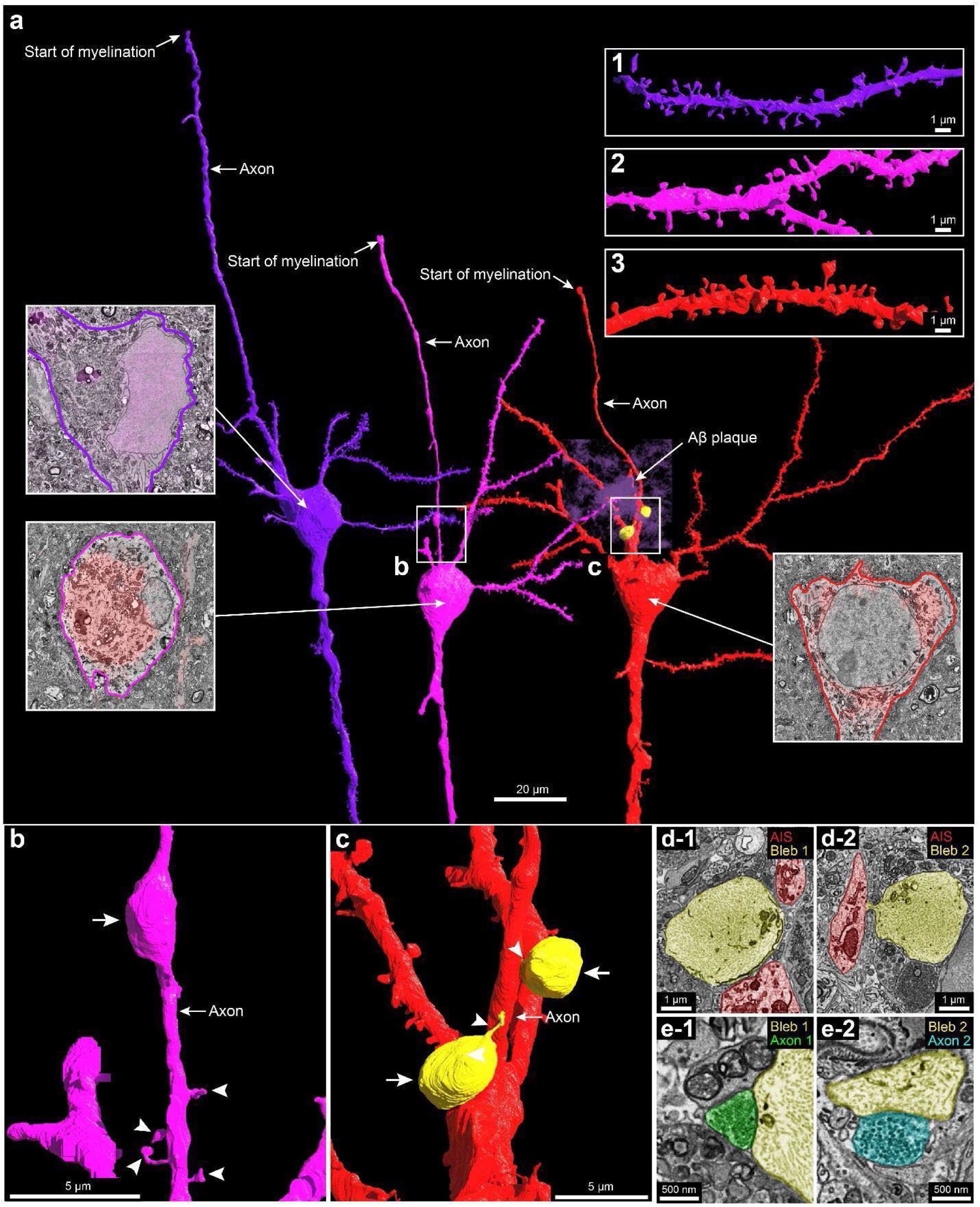
3D reconstruction of a neuron with intracellular Aβ and two neurons with intracellular pTau. **a**, 3D reconstruction of three pyramidal neurons. The insets pointing to the cell bodies show the labeling of Aβ or pTau by nanobody probes. Insets 1 to 3 show segments of spiny dendrites of Neuron 1 to 3. **b** shows the AIS of Neuron 2 as labeled in **a**. White arrow, the swelling filled with pTau. White arrow heads, spines on AIS. **c** shows the AIS of Neuron 3 as labeled in **a**. White arrows, two pTau positive blebs connected to the AIS. White arrows, thin stalks connecting the blebs to the AIS. **d-1** and **d-2**, 2D segmentations of the two pTau positive blebs filled with straight filaments. **e-1** and **e-2** show that each of the blebs was adjacent to an axon profile filled with synaptic vesicles.

### Analysis of synapse nearby an Aβ plaque reveals alterations in synapse density and size

As another proof-of-principle demonstration of the robustness and utility of our dataset, we applied the synapse prediction by supervised learning that has high precision and recall to investigate synaptic alterations in proximity to an Aβ plaque—an area of critical interest given the central role of synaptic pathology in the etiology, manifestation, and progression of AD (Tzioras et al. 2023; Koffie, Hyman, and Spires-Jones 2011). We randomly selected nine 6 x 6 x 6 µm regions predominantly comprising neuropil. These regions included three representative areas located at distances of 10 µm, 20 µm, and 30 µm from the plaque center (Sup. Figure 9).

Synapse detection on all nine boxes (Figure 10 a-1 to a-3, b-1 to b-3, c-1 to c-3) showed that a decrease in synapse density as regions progressively closer to the plaque: synapse density was 0.181 ± 0.068 per µm³ at 10 µm, 0.653 ± 0.025 per µm³ at 20 µm, 1.216 ± 0.174 per µm³ at 30 µm (Figure 10 d). There was a statistically significant trend as the distance to the plaques dropped from 30 µm to 10 µm (two-tailed, unpaired t-test. 10 µm vs. 20 µm, p = 0.0004; 20 µm vs. 30 µm, p = 0.0051; 10 µm vs. 30 µm, p = 0.0007).

**Figure 10.**
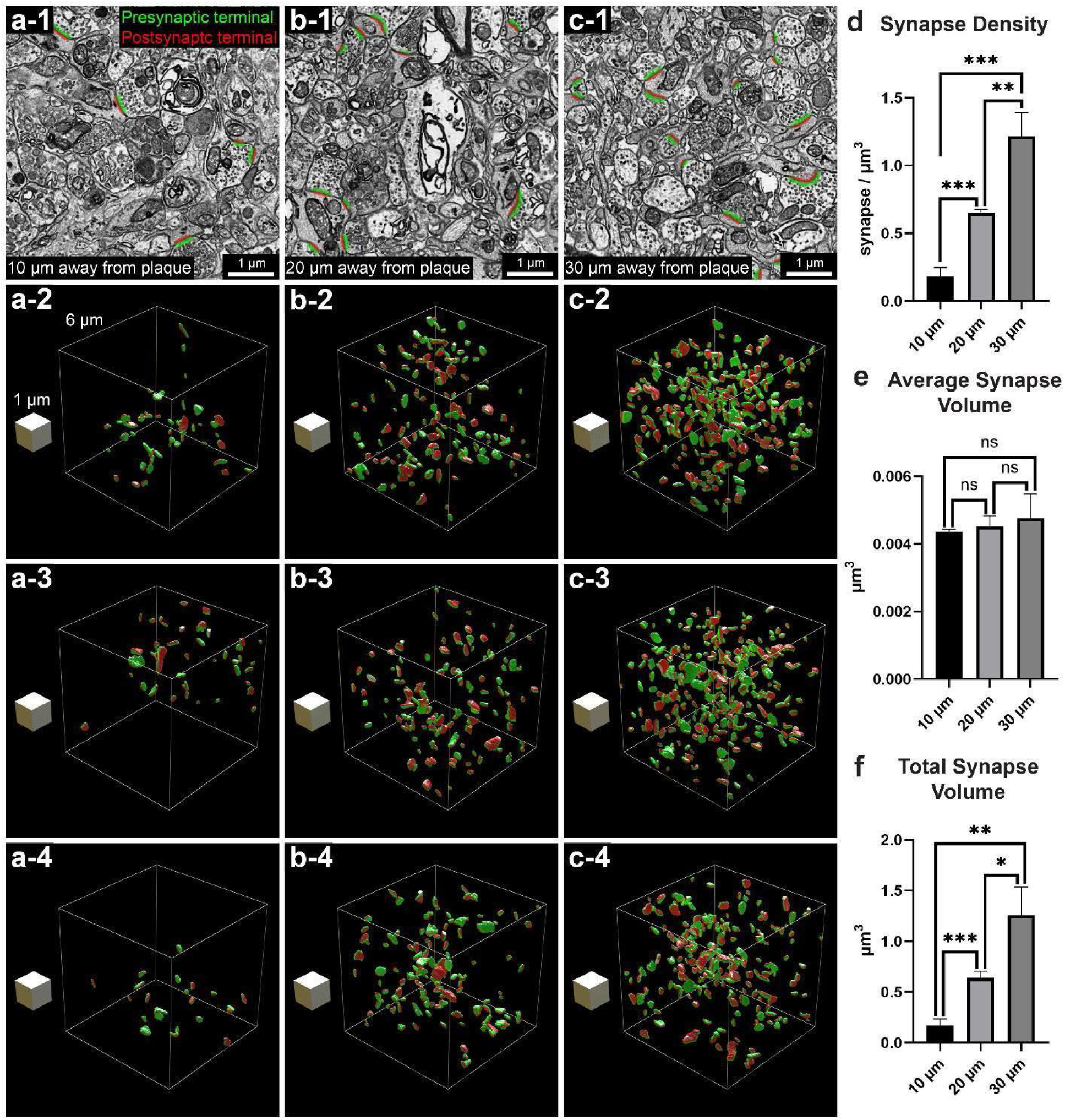
Changes in synapse density and total synapse volume with relationship to the distance from an extracellular Aβ plaque. **a-1**, A representative image slice with detected synapses from the box at 10 µm away from the center of an Aβ plaque. **a-2** to **a-4**, 3D rendering of all detected synapses in the box as in **a-1** with the dimensions of 6 µm x 6 µm x 6 µm at 10 µm away from the center of an Aβ plaque. White block is of the dimension of 1 µm x 1 µm x 1 µm. **b-1**, A representative image slice with detected synapses from the box at 20 µm away from the center of an Aβ plaque. The image slice is of the dimension of 6 µm x 6 µm. **b-2** to **b-4**, 3D rendering of all detected synapses in the box as in **b-1** with the dimensions of 6 µm x 6 µm x 6 µm at 20 µm away from the center of an Aβ plaque. White block is of the dimension of 1 µm x 1 µm x 1 µm. **c-1**, A representative image slice with detected synapses from the box at 30 µm away from the center of an Aβ plaque. The image slice is of the dimension of 6 µm x 6 µm. **c-2** to **c-4**, 3D rendering of all detected synapses in the box as in **c-1** with the dimensions of 6 µm x 6 µm x 6 µm at 30 µm away from the center of an Aβ plaque. White block is of the dimension of 1 µm x 1 µm x 1 µm. **d**, Synapse density, **e**, Average synapse volume, **f**, Total synapse volume were measured for each of the three box at 10 µm away from the plaque, the three box at 20 µm away from the plaque, and the three box at 30 µm away from the plaque. Two-tailed, unpaired t-test was performed. Synapse density, 10 µm vs. 20 µm, p = 0.0004, t=11.29, df=4; 20 µm vs. 30 µm, p = 0.0051, t=5.553, df=4; 10 µm vs. 30 µm, p = 0.0007, t=9.597, df=4. Average synapse volume, 10 µm vs. 20 µm, p = 0.4356, t=0.8655, df=4; 20 µm vs. 30 µm, p = 0.6104, t=0.5519, df=4; 10 µm vs. 30 µm, p = 0.3864, t=0.9713, df=4. Total synapse volume, 10 µm vs. 20 µm, 0.0009, t=8.789, df=4; 20 µm vs. 30 µm, p = 0.0210, t=3.690, df=4; 10 µm vs. 30 µm, p = 0.0029, t=6.489, df=4.

The average synapse volume (pre-synaptic side) was 0.00436 ± 0.00007 µm³ at 10 µm, 0.00452 ± 0.00030 µm³ at 20 µm, 0.00476 ± 0.00071 µm³ at 30 µm (Figure 10 e). There was no statistically significant change in average synapse volume as the distance to the plaques dropped from 30 µm to 10 µm (two-tailed, unpaired t-test. 10 µm vs. 20 µm, p = 0.4356; 20 µm vs. 30 µm, p = 0.6104; 10 µm vs. 30 µm, p = 0.3864).

The total synaptic volume (pre-synaptic side) for each was 0.170 ± 0.065 µm³ at 10 µm, 0.638 ± 0.066 µm³ at 20 µm, 1.255 ± 0.282 µm³ at 30 µm (Figure 10 f). There was a statistically significant decrease in total synaptic volume as the distance to the plaques dropped from 30 µm to 10 µm (two-tailed, unpaired t-test. 10 µm vs. 20 µm, p = 0.0009; 20 µm vs. 30 µm, p = 0.0210; 10 µm vs. 30 µm, p = 0.0029).

To investigate whether smaller synapses are more susceptible to elimination, suggested by previous studies (John and Reddy 2021; Meftah and Gan 2023)—we combined synapse volume data (pre-synaptic + post-synaptic) from the three boxes at each distance and analyzed the frequency distributions (Sup. Figure 10). We performed two-sample Kolmogorov–Smirnov tests and found there was no significant difference between any two of the distributions (10 μm vs. 20 μm: p = 0.8498; 20 μm vs. 30 μm: p = 0.1231; 10 μm vs. 30 μm: p = 0.09247). This result suggests that there was synapse elimination closer to the plaque, but the elimination did not preferentially target smaller nor larger synapses. Synapses of all volumes were possibly eliminated equally when closer to the plaque. To further test whether one of the pre-synaptic site or post-synaptic is affected more, we plotted the frequency distributions of pre- or post-synaptic volumes (pre-synaptic, Sup. Figure 11 a; post-synaptic, Sup. Figure 11 b) for all nine boxes. We found that the pre-synaptic volumes of all boxes were slightly larger than the post-synaptic volumes (Sup. Figure 11 c).

## Conclusions and Discussion

This work introduces a volumetric correlated light and electron microscopy (vCLEM) technique using fluorescent nanobody immunolabeling to localize Alzheimer’s disease (AD)-related molecules, including phosphorylated tau (pTau), amyloid-β (Aβ), and CD11b, in a large-scale volume EM dataset from an AD mouse model. By preserving ultrastructure through detergent-free labeling, this method addresses a major limitation of conventional volume EM-based connectomics, which lacks molecular specificity, and overcomes the challenges posed by traditional molecular labeling techniques that often compromise sample integrity. The resulting dataset enables precise localization of pathological proteins within their ultrastructural context, allowing for integrated analyses at molecular, cellular, and circuit levels across a significantly larger volume than previous vEM datasets.

<u>The publicly accessible dataset</u>, featuring advanced machine-learning-based segmentations of neurons, dendrites, axons, glial processes, and synapses, offers a valuable resource for studying AD pathology. Researchers can pinpoint cellular and subcellular structures where Aβ and pTau localize, reconstruct affected cells at nanometer resolution, and investigate how these pathological markers impact synaptic connectivity and neuronal networks. This dataset lowers the entry barrier for researchers less familiar with volume EM or segmentation tools, fostering broader participation in connectomics-driven AD research.

Following the above framework, as a proof-of-principle, we used our dataset to identify novel pTau-associated abnormalities at the axon initial segment (AIS), including enlarged blebs filled with straight filaments and pTau-positive swellings containing lipofuscin granules (Figure 6 g-1, g-2). These findings highlight the complex landscape of pTau pathology, where certain aggregates lack visible neurofibrillary tangle-like features in EM micrographs. We also observed spine-like structures at the AIS with thin stalks (Figure 9 c; less than 1 µm in diameter), potentially prone to detachment, raising the intriguing hypothesis that these structures may contribute to exosome formation, a mechanism implicated in pTau release in AD (Saman et al. 2012; Miyoshi et al. 2021). These abnormalities suggest disrupted molecular mechanisms at the AIS, likely involving ankyrin-G (Sobotzik et al. 2009; Zempel et al. 2017) or diffusion barriers affected by pTau or Aβ (X. Li et al. 2011), leading to tau missorting and synaptic dysfunction (Blazquez-Llorca et al. 2011). The AIS’s established role as a barrier for regulating tau trafficking underscores the potential of these disruptions to impact synaptic transmission significantly. For Aβ, in addition to extracellular deposits in plaques, we identified neurons with intraneuronal Aβ deposits exhibiting severe morphological abnormalities, including tortuous dendrites, shrunken cell bodies, and degenerating organelles. These features indicate profound cellular dysfunction and are consistent with Aβ’s known neurotoxicity. Interestingly, 3D reconstructions enabled by machine-learning segmentation provided a detailed view of these affected neurons and revealed novel morphologies not easily captured by prior FIB-SEM and ATUM-SEM datasets, which were limited to smaller subvolumes (Montero-Crespo et al. 2021; Domínguez-Álvaro et al. 2018; Jiang et al. 2022; Pang et al. 2022; Blazquez-Llorca et al. 2013).

Our synapse analysis further revealed significant alterations near Aβ plaques. Synapse density declined sharply with proximity to plaques, reflecting Aβ’s synaptotoxic effects (Hampel et al. 2021; Koffie et al. 2009; Urbanc et al. 2002) and glial-mediated mechanisms (Chung et al. 2015; Rajendran and Paolicelli 2018; Vilalta and Brown 2018). Reactive glial processes from microglia and astrocytes were notably abundant near plaques (Figure 8 b, also see Sup. Figure 11 for astrocytic processes), supporting their role in synaptic pruning and dysfunction. While synapse density decreased, the average synapse volume remained constant, resulting in a reduction in total synapse volume near plaques. This challenges earlier findings suggesting compensatory synaptic enlargement with declining synapse density (Scheff and Price 2001; Neuman et al. 2015; Androuin et al. 2018; Scheff and Price 2003). Instead, our results align with more recent volume EM studies showing no evidence of synaptic enlargement (Martínez-Serra et al. 2022). Future classification of synapses by type (e.g., excitatory vs. inhibitory) or location (e.g., somatic vs. dendritic) could clarify whether specific subpopulations respond differently to Aβ toxicity.

While the vCLEM approach provides transformative insights, it has limitations. We focused on two canonical markers—fibrillar Aβ40/42 and pTau phosphorylated at specific sites—without addressing other molecular forms known to elicit more severe toxicity. Additionally, the limited availability of nanobody probes constrains the range of molecules that can be studied. The resolution gap between confocal fluorescence microscopy and EM may introduce slight inaccuracies in registering molecular signals onto EM micrographs. Furthermore, findings from this single-sample study require validation in additional datasets, including other animal models and human tissues, to ensure broader applicability.

A key future direction involves expanding the repertoire of fluorescent nanobody probes to target a broader range of pathological molecules. For instance, probes could be developed for specific Aβ species (e.g., Aβ1-40 and Aβ1-42) and conformations (e.g., oligomers, protofibrils, and fibrils) (Hampel et al. 2021), as well as tau isoforms or site-specific phosphorylations (Arendt, Stieler, and Holzer 2016). These advancements would enable more detailed analyses of molecular variants with distinct toxicities affecting neurons and synapses. Furthermore, nanobody probes for other neurodegeneration-related molecules, such as α-synuclein (Twohig and Nielsen 2019), TDP-43 (transactive response DNA-binding protein of 42 kDa) (Wilson et al. 2011), PLD3 (Nackenoff et al. 2021; Hooli et al. 2015), and SORL1 (sortilin-related receptor 1), which plays a crucial role in neuronal endosome trafficking (Yin, Yu, and Tan 2015; Knupp et al. 2020), as well as huntingtin (Bates et al. 2015), could significantly broaden the utility of this technique. These additions would expand its application beyond Alzheimer’s disease to other forms of dementia and neurodegenerative disorders. Many nanobodies for these targets already exist (Danis et al. 2022; De Genst et al. 2010; Habicht et al. 2007; Lafaye et al. 2009) and could be easily adapted into fluorescent probes. Beyond pathological proteins, this approach can also be extended to cell-type-specific markers, allowing researchers to investigate how toxic molecules impact distinct cell populations. For example, nanobody or single-chain variable fragment (scFv)-based probes could target markers for glial cells or inhibitory neuron subtypes, providing insights into cell-type-specific vulnerabilities and mechanisms in Alzheimer’s disease and other neurodegenerative conditions.

Recent large-scale proteomic and transcriptomic studies (Johnson et al. 2022, 2020; Magistri et al. 2015; Mathys et al. 2019) have showed alterations at the levels of protein and gene expression in AD. To localize these new molecular abnormalities in connectomics datasets, a large collection of nanobody probes that target common molecular markers will be needed. Multiple nanobody databases are now available where thousands of nanobody sequences are being deposited (Deszyński et al. 2021; Wilton et al. 2018). Researchers can query sequences of nanobodies that target the molecular abnormalities of interest, and then, as we have done, convert these nanobodies into fluorescent probes. If no nanobody for a desired target is available, there are efficient ways to generate new nanobodies from immune libraries (Muyldermans 2021) or synthetic libraries (McMahon et al., 2018; Zimmermann et al., 2020)(McMahon et al. 2018; Zimmermann et al. 2020).

With a growing collection of nanobody probes, integrating vCLEM with super-resolution microscopy (Hoffman et al., n.d.; Tuijtel et al. 2019) could enhance the precision of molecular target localization within EM micrographs. Combining vCLEM with advanced multiplex imaging techniques, such as super-multicolor fluorescence imaging enabled by spectral unmixing (Seo et al. 2022) or multiplexed Raman vibrational imaging (Wei et al. 2017), could allow for the simultaneous labeling of numerous markers in a single sample. This would unlock unprecedented opportunities for integrative molecular and structural studies, enabling a deeper understanding of complex pathological processes.

Notably, this technique is well-suited for application to human samples, as it does not rely on transgenic models and can directly target molecules in human tissues using nanobody probes. The nanobody probes tested in this study (A2 and R3VQ) have already been shown to effectively detect their targets in brain samples from AD patients (T. Li et al. 2016). By integrating this approach with recently developed methods for processing large surgical brain samples (Karlupia et al. 2023; Loomba et al. 2022), vCLEM offers a powerful tool for investigating AD connectomics directly in human tissues, providing critical insights into disease mechanisms and pathology.

This study highlights the transformative potential of vCLEM for bridging molecular and structural data in large-scale volume EM datasets, offering a publicly accessible resource to advance integrative research into AD and other neurodegenerative diseases.

## Material and Methods

### Animals

Animals used in the study were female one-year old 3xTg (B6;129-Tg(APPSwe,tauP301L)1Lfa Psen1^tm1Mpm/Mmjax^) mice (Jackson Laboratory) and female one-year old B6129SF2/J mice (as age-matched control animal, Jackson Laboratory). All experiments using animals were conducted according to US National Institutes of Health guidelines and approved by the Committee on Animal Care at Harvard University.

### Nanobody immuno-probe production

A signal peptide (MDWTWRILFLVAAATGAHS) was added to the N’-terminus of the nanobody. A sortase tag (SLPETGG) and a 6 x His tag was added to the C’-terminus. The amino acid sequence was reverse translated into a DNA sequence with codon optimization for human cells. The DNA sequence was synthesized and cloned into pcDNA 3.1 vectors.

Nanobody expression was performed with Expi 293 cells (ThermoFisher). Detailed expression, purification, and dye conjugation protocols can be found in Sup. File.

### Perfusion and fixation

The mouse was anesthetized by isoflurane until there was no toe-pinch reflex. Mice were then transcardially perfused with aCSF (125 mM NaCl, 26 mM NaHCO_3_, 1.25 mM NaH_2_PO_4_, 2.5 mM KCl, 26 mM glucose, 1 mM MgCl_2_ and 2 mM CaCl_2_ (all chemicals from Sigma-Aldrich) at the flow rate of 10 ml/min for 2 min to remove blood, followed with 4% paraformaldehyde (Electron Microscopy Sciences), 0.1% glutaraldehyde (Electron Microscopy Sciences) in 1 x PBS for 3 min for fixation. Brains were dissected and then post-fixed in the same fixative on a rotator overnight at 4°C. Brains were sectioned into 50-μm or 120-μm coronal sections using a Leica VT1000 S vibratome and stored in the same fixative at 4°C.

### Immunofluorescence

Detergent-free immunofluorescence labeling was performed with nanobody probes (see Sup. Table 1 for the nanobody probes and the final concentration of each nanobody probe used in this study). The detailed protocol was described in (Han et al. 2024). For the 120-µm coronal sections used for vCLEM, the section was first washed with 1 x PBS for 3 x 10 min, and then blocked in glycine blocking buffer (0.1 M glycine, 0.05% NaN_3_ in 1 x PBS) for 1 h on a rotator at 4 °C. The labeling solution was prepared by diluting nanobody-dye conjugates in glycine blocking buffer (see Sup. Table 1 for the final concentration used). The labeling solution and any remaining steps were protected from light. The section was incubated with the labeling solution on a rotator at 4 °C for three to seven days. The 120-µm sections for Figure 2 were first incubated with the labeling solution with only the anti-pTau nanobody for three days, and then incubated with the labeling solution with the anti-Aβ and the anti-CD11b nanobodies for another three days. The 120-µm section for Figure 3 was incubated with the labeling solution with all four nanobodies for seven days. After the incubation, the section was washed with 1 x PBS for 3 x 10 min, and then stained with Hoechst 33342 (Invitrogen, diluted 1:5000 in 1 x PBS) for 1 h on a rotator at 4 °C. The section was washed with 1 x PBS for 3 x 10 min, and then mounted onto glass slide (VWR).

To validate the labeling of the nanobody A2, double immunofluorescence labeling with detergent was performed on 50-µm sections with the anti-pTau mAb AT8 (ThermoFisher) plus the anti-mouse IgG H + L secondary antibody conjugated with Alexa Fluor 488 (ThermoFisher), and A2 conjugated with Alexa Fluor 594 (see Sup. Figure 1 a) (see Sup. Table 2 and 3 for the final concentrations used for each probe). To validate the labeling of the nanobody R3VQ, double immunofluorescence labeling with detergent was performed with the anti-Aβ mAb 4G8 conjugated with Alexa Fluor 647 (BioLegend) and R3VQ conjugated with 5-TAMRA (see Sup. Figure 2) (see Sup. Table 2 and 3 for the final concentrations used for each probe). The detailed protocol is available on the NeuroMab website (“Welcome to NeuroMab!,” n.d.). In brief, 50-µm coronal sections were first washed with 1 x PBS for 3 x 10 min, and then blocked in vehicle (10% normal goat serum, 0.3% Triton X-100 in 1 x PBS) overnight on a rotator at 4 °C. Subsequent steps were protected from light. For the experiment with AT8 and A2, the sections were then incubated with primary antibody solution plus nanobody probes overnight on a rotator at 4 °C. After the incubation, sections were washed with vehicle for 3 x 10 min, and then incubated with the secondary antibody solution for 1 hour on a rotator at 4 °C. For the experiment with 4G8 and R3VQ, the sections were then incubated with primary antibody solution plus nanobody probes for 3 days on a rotator at 4 °C. After the incubation, sections were washed with 1 x PBS for 3 x 10 min, and then stained with Hoechst (diluted 1:5000 in 1 x PBS) for 1 h on a rotator at 4 °C. Sections were washed with 1 x PBS for 3 x 10 min, and then mounted onto glass slides.

### Fluorescence confocal microscopy

Fluorescence confocal microscopy was performed as described in (Han et al. 2024). In brief, 50-µm sections were mounted in Vectashield H-1000 (Vector Laboratories) with a #1 coverslip (Electron Microscopy Sciences) on top sealed with clear nail polish. 120-µm sections (for vCLEM) were mounted in 1 x PBS inside 120-um spacer (Invitrogen) with a #1 coverslip on top sealed with the spacer. Sections were imaged with a Zeiss LSM 880 confocal laser scanning microscope equipped with either a 20x/0.8 NA air-objective or a 40x/1.1 NA water immersion objective. The 120-µm section for vCLEM was imaged with a 40x/1.1 NA water immersion objective. Acquisition of double or triple color fluorescent images was done with appropriate band pass filters for the specific fluorescent dyes to avoid crosstalk.

The brightness, contrast, and gamma of all fluorescent images were adjusted. Fluorescence image volumes were projected to a single plane by maximum intensity for visualization in 2D.

### EM preparation

Sample preparation for electron microscopy was performed as described in (Han et al. 2024). After the 120-µm sections was imaged, it was transferred into 3.7 ml shell vials with 1 ml secondary fixative (2% PFA, 2,5% glutaraldehyde in 0.15M sodium cacodylate buffer with 4mM Ca^2+^ and 0.4 mM Mg^2+^) and incubated for at least one week on a rotator at 4 °C. A modified ROTO (Reduced Osmium-Thiocarbohydrazide-Osmium) protocol was used to stain the section. The section was embedded in LX-112 resin.

### X-ray microCT scanning

X-ray micro-computed tomography (μCT) images of the resin-embedded sections were acquired using a Zeiss Xradia 510 Versa system and Zeiss’ Scout and Scan software. Detailed protocol was described in (Han et al. 2024).

### EM imaging

The resin-embedded sections were cut into 30 nm serial ultrathin sections using automated tape-collecting ultramicrotome (ATUM) (Kasthuri et al. 2015). Serial sections were collected onto carbon-coated and plasma-treated Kapton tape. The tape was cut into strips and affixed onto 150 mm silicon wafers (University Wafer). A Zeiss Sigma scanning electron microscope was used to acquire overview images from the serial sections. Detailed protocol was described in (Han et al. 2024).

Prior to acquiring high-resolution images, the serial section sections on wafers were post-stained for 4 min with a 3% lead citrate solution. After staining, the sections were degassed for a minimum of 24 h at 1×10-6 Torr. A Zeiss MultiSEM 505 scanning electron microscope equipped with 61 electron beams was used to acquire high-resolution images from the serial sections. Images were collected using a 1.5-kV landing energy, 4-nm image pixel, and a 400-ns dwell time.

### High-resolution EM image processing

The preparation of the vEM data before it could be segmented/analyzed includes two steps: affine stitching and elastic alignment. Detailed protocol was described in (Han et al. 2024). We excluded 20 slices that had sub-optimal imaging quality (e.g. very thin slices or contains tears and wrinkles) and copied the adjacent sections in the places of these excluded slices. The aligned stack was rendered at full resolution (4 x 4 x 30 nm) and each section was cut into 4k x 4k .png tiles, imported into VAST as a .vsvi file, and ingested into Neuroglancer for further analysis.

The initial alignment of the vEM dataset showed several jumps through the z-axis. A second attempt at alignment was performed after discarding four more slices (we also copied the adjacent sections in the places of these discarded slices) that had suboptimal image quality, which successfully eliminated the jumps. The 3D reconstruction of Neuron 3, the automatic Aβ plaque detection, and the synapse detection using a U-Net classifier was performed on the first version of alignment. The 3D reconstruction of Neuron 3 and the automatic Aβ plaque detection were later transferred to the second version of the alignment based on matching coordinates. The other reconstructions and the synapse detection by PyTorch were performed on the second version of the alignment. The publicly accessible vCLEM dataset shows the second version of the alignment.

### Co-registration of fluorescence and EM volumes

The co-registration was performed using the BigWarp plugin in FIJI. Detailed description can be found in (Han et al. 2024).

### Automatic segmentation

We segmented a central cutout of the dataset using flood-filling networks (Michał Januszewski et al. 2018). The FFN segmentation model was trained at 16 x 16 x 33 nm nominal resolution on the H01 dataset (Shapson-Coe et al. 2024), and run here on CLAHE intensity normalized data (Zuiderveld 1994) downsampled to 16 x 16 x 30 nm. Direct application of the H01-trained FFN to the AD dataset did not perform well, particularly on thinner objects. Therefore we used Segmentation Enhanced CycleGANs (Michal Januszewski and Jain 2019) to modify the visual appearance (photometrics, noise statistics, staining, etc.) of the AD dataset at 16 x 16 x 30 nm to mimic H01, prior to running the FFN. We trained the SECGAN style transfer on a central cutout of the AD dataset totaling 240 x 160 x 15 um and a cutout of H01 totaling 1600 x 760 x 130 um for 8 million steps at batch size 1. The SECGAN generator architecture was an 8-layer fully convolutional stack with residual connections and no downsampling, and the discriminator was a ResNet18 (He et al., n.d.).

2D membrane detection at 4 x 4 x 30 nm resolution was performed on the vEM dataset using a method developed in our lab. A description of this approach was given in (Karlupia et al. 2023; Meirovitch et al. 2018; Pavarino et al. 2023). In brief, an algorithm (Pavarino et al. 2023) pre-trained on a mouse cerebellar dataset was used to generate 2D membrane detection. 2D membrane detection was use for filtering false positive in synapse prediction.

### Synapse detection by supervised leaning

We sampled nine EM image subvolumes with dimensions of 6 µm x 6 µm x 6 µm at three distances (10 µm, 20 µm, 30 µm) from the center of an Aβ plaque at the original resolution. We took a semi-automatic approach to annotate all the synapses in these subvolumes. First, one human annotator labeled all the pre- and post-synaptic appositions in 1/4 of one of the nine EM image subvolumes. Then, we trained a synapse detection model (Parag et al. 2019) using the manual annotations with the PyTorch Connectomics deep learning package. To refine the model, we tested it on all slices of this subvolume, manually proofread its results, and fine-tuned the model with all the annotations, which led to a desirable detection accuracy and recall result. Lastly, we used the improved model to detect synapses in the entire dataset and identified 3.5 million synapses..

### Synapse detection by transfer learning

The original synapse annotation model was a UNet (3-down and 3-up modules) trained at 8 x 8 x 33 nm nominal resolution on the H01 dataset (Shapson-Coe et al. 2024). For the initial synapse predictions, we used the SECGAN model trained for 683k steps to style-transfer an ROI of the AD data to mimic H01, and then ran a synapse model that was trained for 786k steps on H01. This produced 0.255 million synapse masks, with low rates of precision and recall. We then used the model-inferred synapse masks as automated ground truth to train a new UNet from scratch directly on the AD data (8 x 8 x 30 nm resolution, without SECGAN style transfer, but with CLAHE). We trained this model for 38M steps and biased it against false negatives by weighting positively labeled pre- and post-synaptic voxels in the training data by 2x. The new synapse annotation model produced 2.1 million raw synapse masks, which resulted in 0.732 million total synapses after filtering for synapses above a 100 voxel volume threshold, removing any pairs that had identical pre- and post-synaptic parents (i.e. self-synapses) and masking out synaptic sites residing on blood vessels, somas, or myelin. We estimated the final precision and recall as 46% and 52% based on evaluation against the manualy annotated subvolume in the first method of synapse detection.

### Automatic Aβ plaque segmentation

We optimized a DeepLab-v3 semantic segmentation model (L.-C. Chen et al. 2018) on a cropped stack of images with manually identified Aβ plaque. The model was pre-trained on another vEM dataset from the Alzheimer’s disease mouse model Tg2576 (Dr. Olga Morozova, personal communication) to decrease the requirements of manually annotated training data in the target volume. We also oversample patches near the Aβ plaque border during training to improve segmentation quality as the mask structures in those patches are more challenging. The training and inference are implemented with the open-source PyTorch Connectomics codebase.

### Statistical analysis

Two-tailed, unpaired t-tests on synaptic density, synaptic volume and total synaptic volume of the three categories defined in the section of synapse detection (The PyTorch method) were performed in Prism-GraphPad. Two-sample Kolmogorov–Smirnov test was performed using Python.

## Acknowledgement

We thank the Harvard Center for Biological Imaging (RRID:SCR_018673) for infrastructure and support, the Biopolymers and Proteomics Core Facility at the Koch Institute at MIT for processing the peptide-fluorescent dye conjugates, the Bauer Core Facility at Harvard for infrastructure and support, Dr. Olga Morozova for helpful discussion on the Alzheimer’s disease mouse model and immunolabeling of Aβ plaques, and Dr. Marta Montero-Crespo for helpful discussion on synapse annotation. This work was supported by NIH grants U19 NS104653, UG3 MH123386, and P50 MH094271 (J. W. L.); NSF grants NSF-CAREER-2239688 (D. W.), NCS-FO-2124179, and NCS-FO-1835231 (H. Pfister.), NLM T15LM007092-31 (M. S.); Office of Naval Research grant N00014-20-1-2828 (J. W. L). X. H. was supported by the Edward R. and Anne G. Lefler predoctoral fellowship from the Lefler Center for Neurodegenerative Disorders at Harvard Medical School and the Simmons Awards from the Harvard Center for Biological Imaging.

## Author Contribution

X. H. and J. W. L conceived this study. Nanobody A2 and R3VQ were originally generated in the labs of P. L., S. B., and B. D.. X. H. generated the fluorescent nanobody probes A2 and R3VQ. H. Ploegh provided the fluorescent anti-CD11b, anti-GFAP, and anti-Ly6C/6G nanobodies. X. H. performed the LM experiments. X. H. and R. S. performed the EM experiments. S. W. performed the imaging processing of the vEM dataset. P. H. L. and V. J. performed 3D segmentation and curated the vCLEM dataset at Neuroglancer. X. H., M. S., and F. A. performed 3D reconstruction of neurons. S. A., B. G., D. W., and X. H. performed synapse prediction through supervised learning.T. B. performed synapse detection through transfer learning. Y. M. performed 2D membrane detection. M. S. performed the filtering of the synapse detection results using a U-Net classifier. Z. L., and H. Pfister performed automatic plaque detection. D. B. provided the reconstruction tool VAST and advice on 3D reconstruction and rendering. Y. W. helped with LM and EM co-registration. X. H. and J. W. L wrote the paper with inputs from H. Ploegh, B. D., D. B., and R. S.

