## Supplementary material for "Mapping Alzheimer’s Molecular Pathologies in Large-Scale Connectomics Data: A Publicly Accessible Correlative Microscopy Resource": Sup. Figure 1

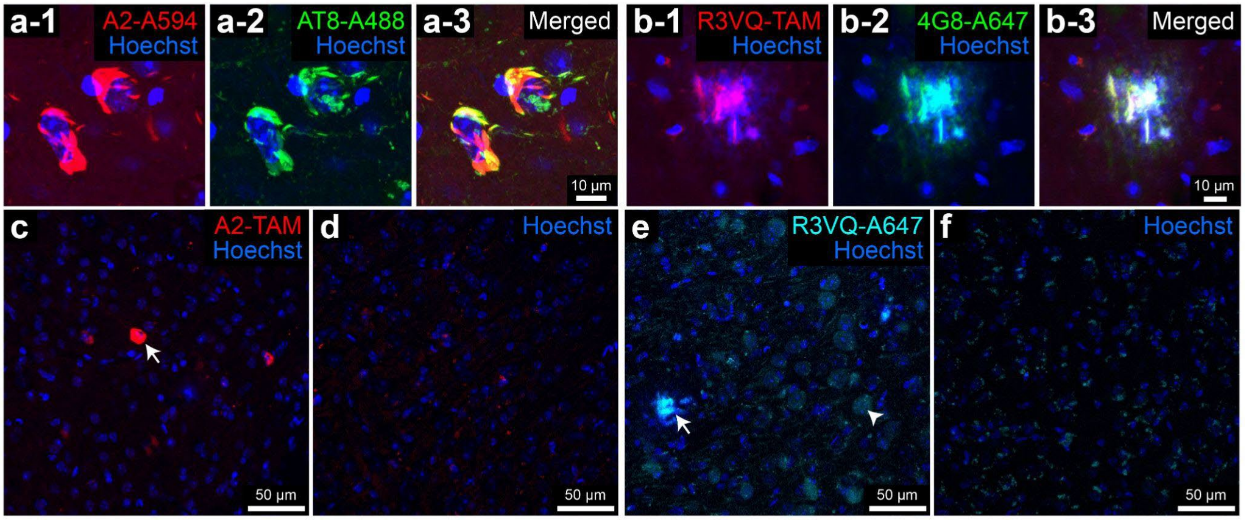


**Sup. Figure 1. Validation of fluorescent nanobody probes for pTau and Aβ.**

**a-1** to **a-3**, Confocal image from the hippocampus of a 3xTg mouse labeled with a pTau-specific nanobody probe (A2) conjugated with Alexa Fluor 594, pTau-specific mAb AT8, and secondary antibodies conjugated with Alexa Fluor 488. **b-1** to **b-3**, Confocal image from the hippocampus of a 3xTg mouse labeled with an Aβ-specific nanobody probe (R3VQ) conjugated with 5-TAMRA and Aβ-specific mAb 4G8 conjugated with Alexa Fluor 647. **c,** Confocal image from the hippocampus of a 3xTg mouse labeled with a pTau-specific nanobody probe (A2) conjugated with 5-TAMRA. The arrow indicates a labeled neuronal cell body. **d**, Confocal image from the hippocampus of an age-matched control mouse labeled with a pTau-specific nanobody probe (A2) conjugated with 5-TAMRA. **e**, Confocal image from the hippocampus of a 3xTg mouse labeled with an Aβ-specific nanobody probe (R3VQ) conjugated with Alexa Fluor 647. The arrow indicates the labeled plaque material. The arrowhead indicates a labeled neuronal cell body. **f**, Confocal image from the hippocampus of an age-matched control mouse labeled with an Aβ-specific nanobody probe (R3VQ) conjugated with Alexa Fluor 647.

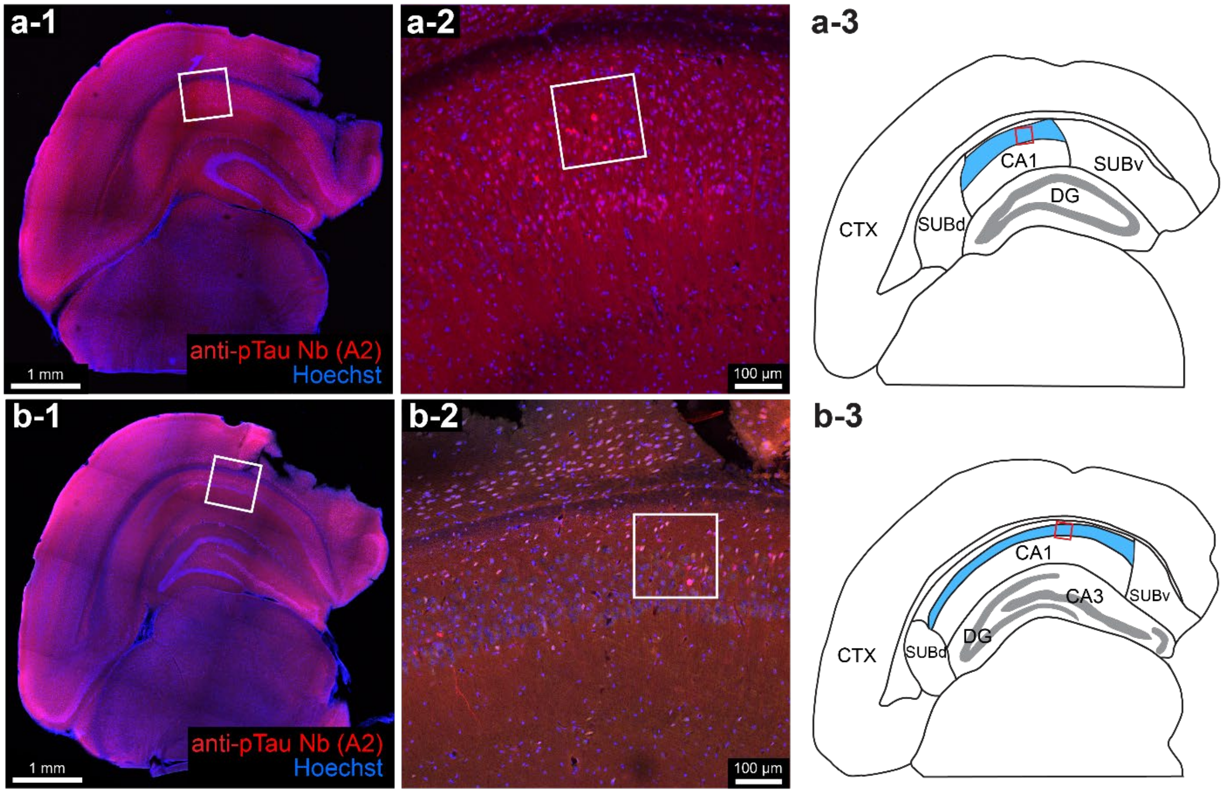


**Sup. Figure 2. Anatomical details of the labeled sections and imaged regions of interest as in Figure 2.**

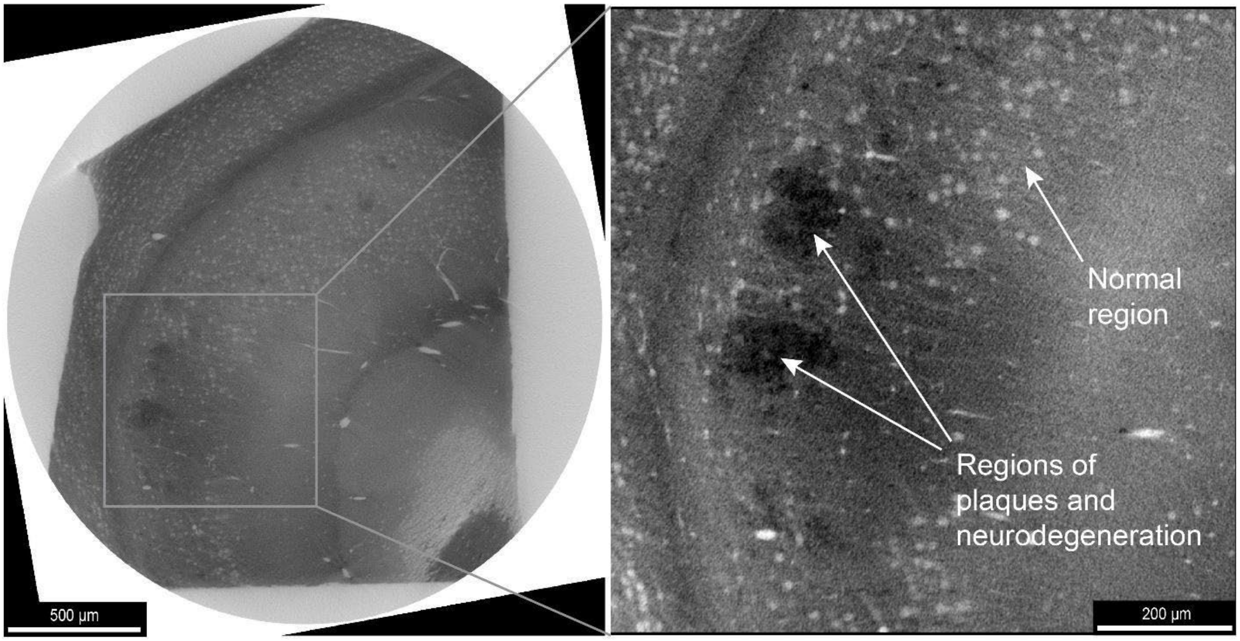

**Sup. Figure 3. An image slice from the X-ray CT (microCT) volume**

Normal regions contain distinguishable cell bodies (light dots), while the regions of plaques and associated neurodegeneration are osmiophilic (darker) and devoid of distinguishable cell bodies.

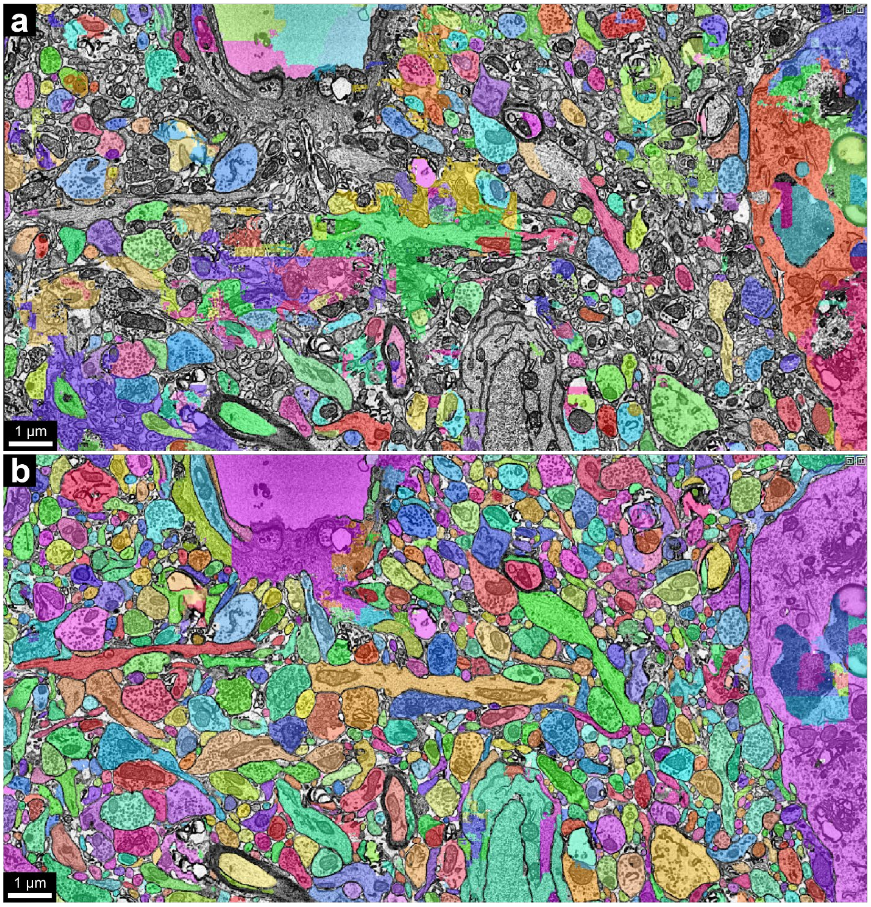


**Sup. Figure 4. Improvement of 3D segmentation by Segmentation Enhanced CycleGANs.**

Comparison between the segmentation (**a**) by directly applying pre-trained FFNs and the segmentation (**b**) adjusted by Segmentation Enhanced CycleGANs.

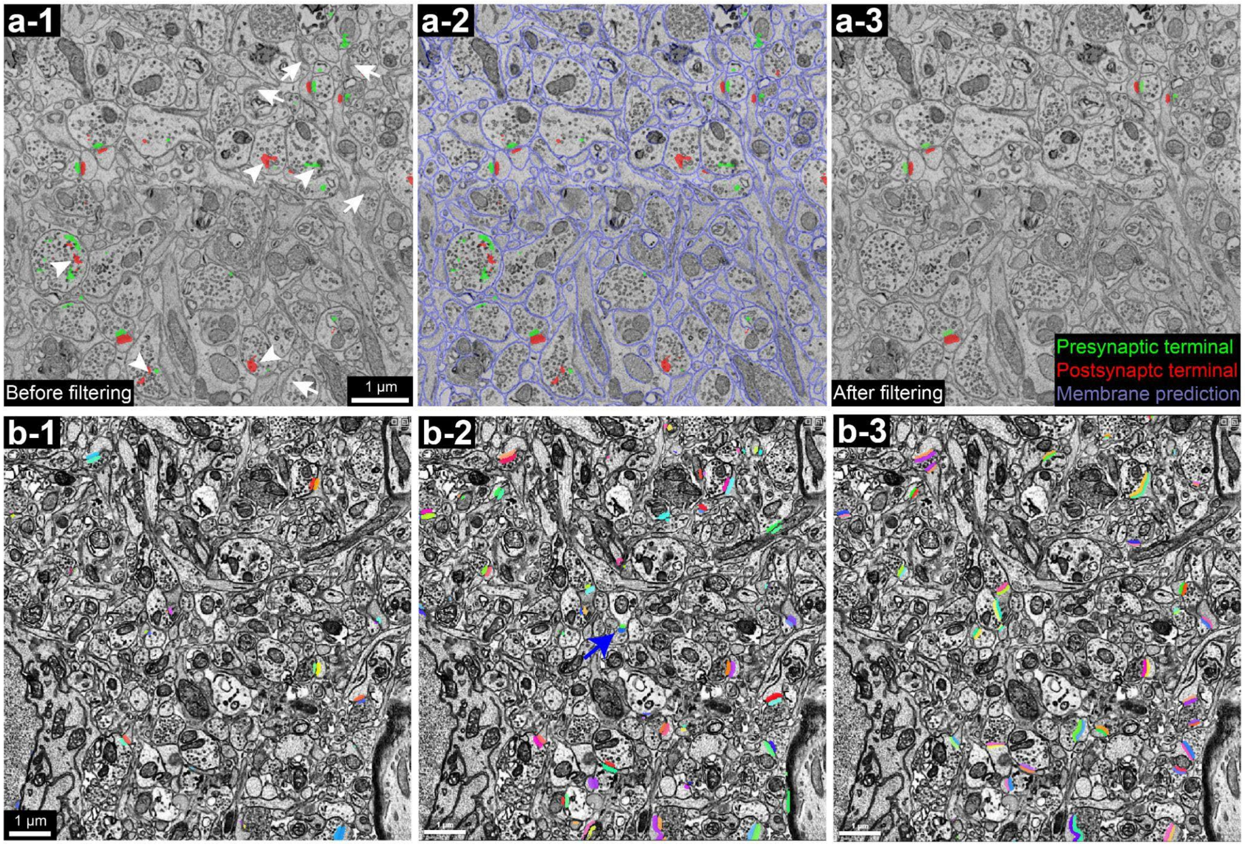


**Sup. Figure 5. Comparison between synapse predictions by the transfer learning and the supervised learning.**

**a-1** to **a-3**, Synapses detected in a region of neuropil by directly running the synapse model trained on the human cortex dataset without transfer learning (**a-1**). By dilating the automatically detected plasma membrane (labeled in purple in **a-2**), we filtered out the false positive detections within neuronal processes (**a-3**). Arrows, synapses that were missed by the detection. Arrowheads, false positive detections within neuronal processes. **b-1** to **b-3**, Comparison between synapse detection results without transfer learning (**b-1**), after the transfer learning (**b-2**), and by the supervised learning (**b-3**). The blue arrow in **b-2** indicates a synapse detected by the transfer learning but missed by the supervised learning.

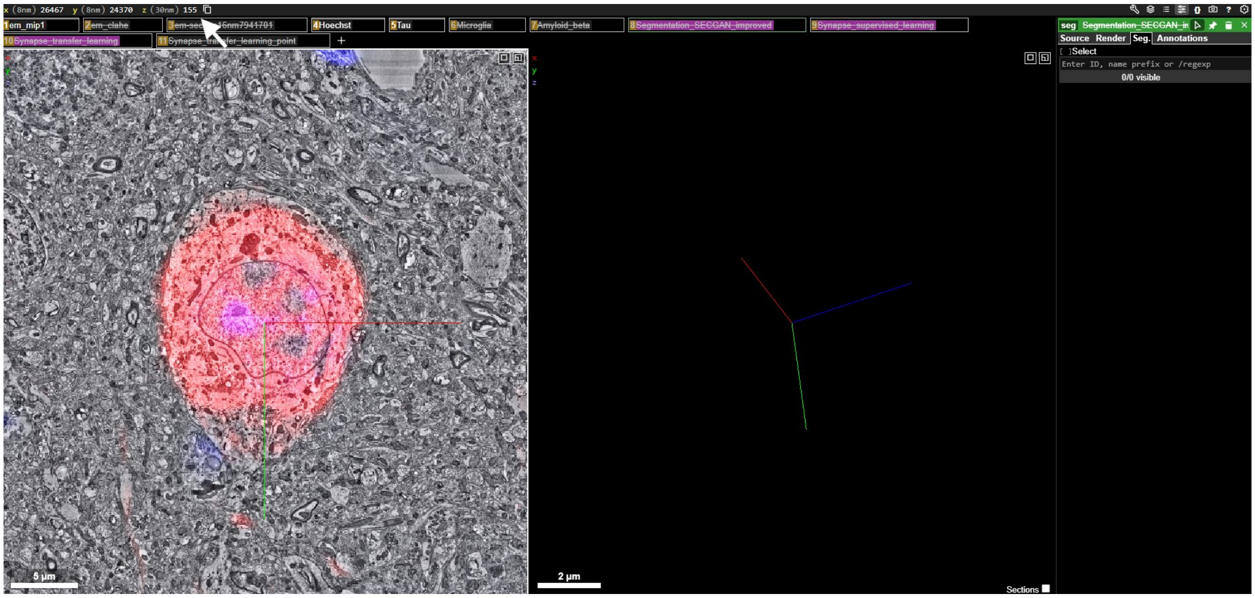


**Sup. Figure 6. Coordinates of an abnormality (labeling pTau in neuronal soma) can be entered into Neuroglancer (arrow) to move to the location.**

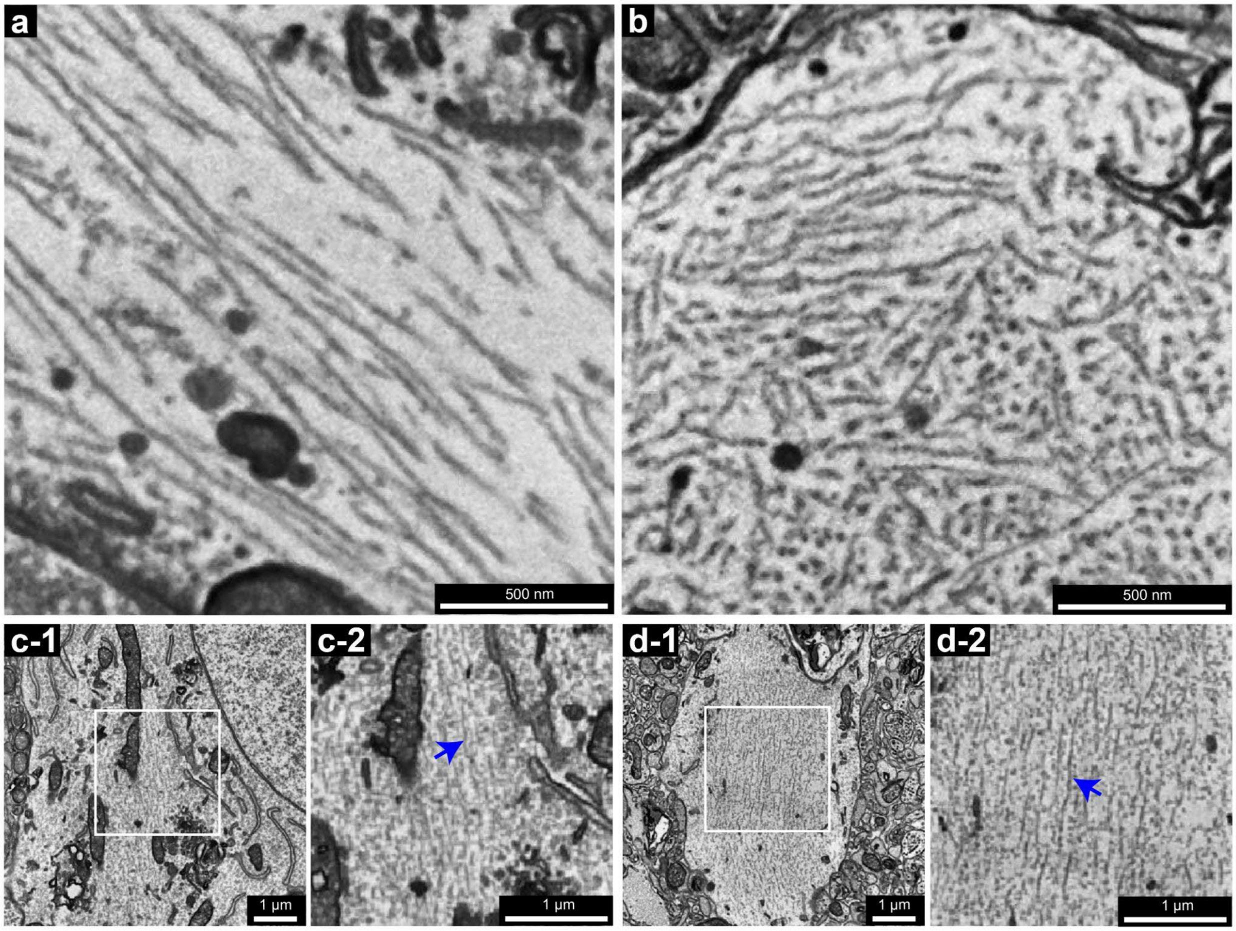

**Sup. Figure 7. Ultrastructural abnormalities related to the anti-pTau nanobody labeling and the comparisons with an unlabeled neuron.**

**a**, high resolution (pixel size = 4 nm) EM micrograph of the straight filament-like structures in Figure 5 a-2. **b**, high resolution (pixel size = 4 nm) EM micrograph of the straight filament-like structures in Figure 5 g-2. **c-1**, a pyramidal neuron that was not labeled by the anti-pTau nanobody, showed short filament-like structures. **c-2**, enlarged inset from **c-1**. The blue arrow shows the short filament-like structures. **d-1**, the same neuron showed short filament-like structures in its apical dendrite. **d-2**, enlarged inset from **d-1**. The blue arrow shows the short filament-like structures.

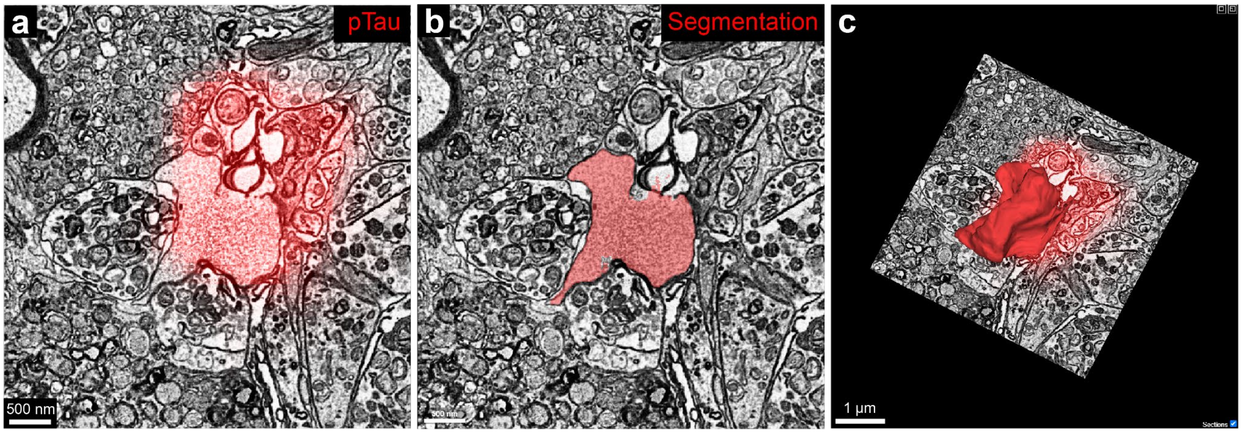


**Sup. Figure 8. An isolated pTau-filled bleb disconnected to any nearby objects.**

**a**, Red fluorescence from the anti-pTau nanobody overlaps with the bleb. **b**, 2D segmentation of the bleb. **c**, 3D reconstruction of the bleb.

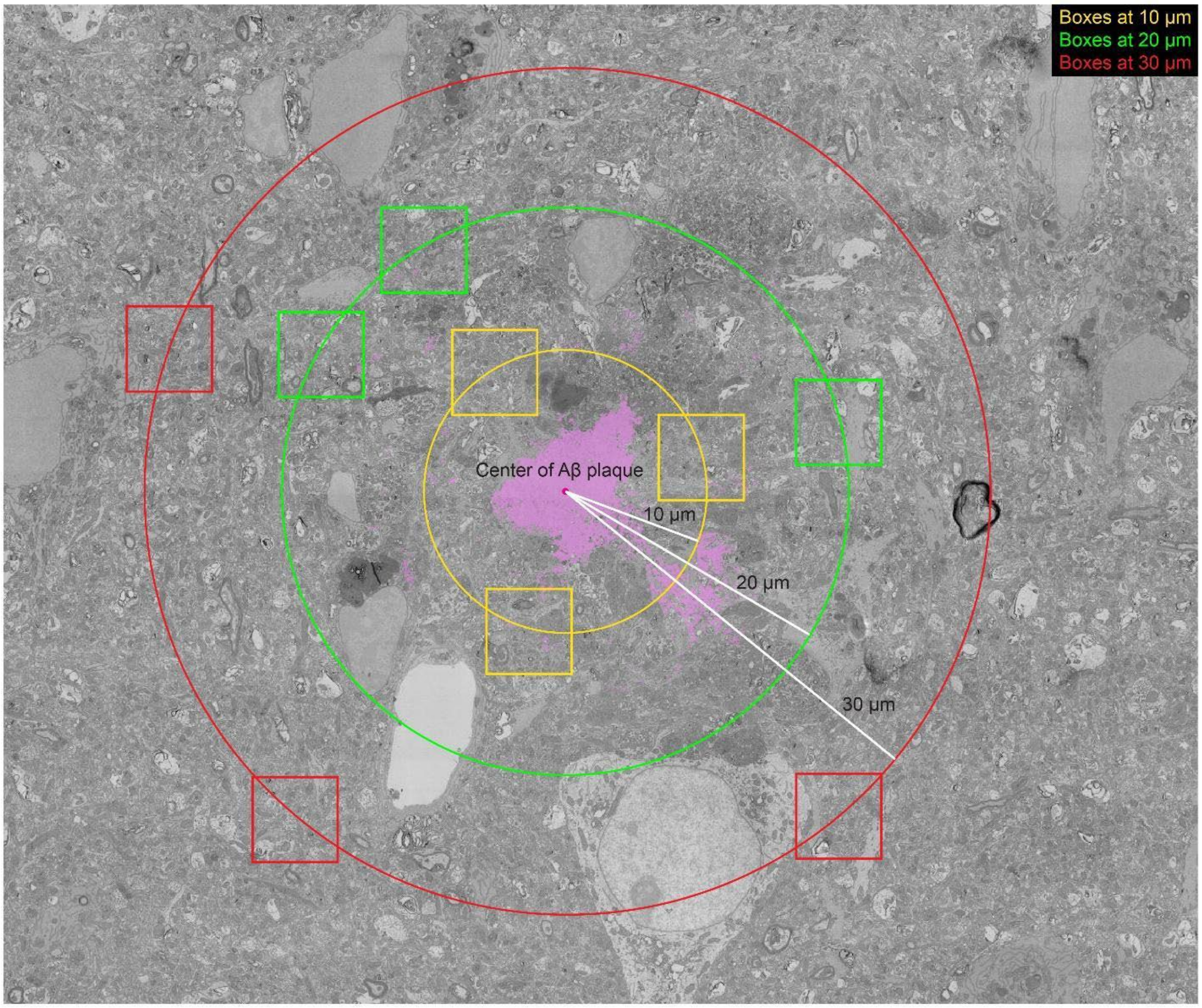


93

94 **Sup. Figure 9. Locations of nine boxes at three distances from the center of a plaque.**

95

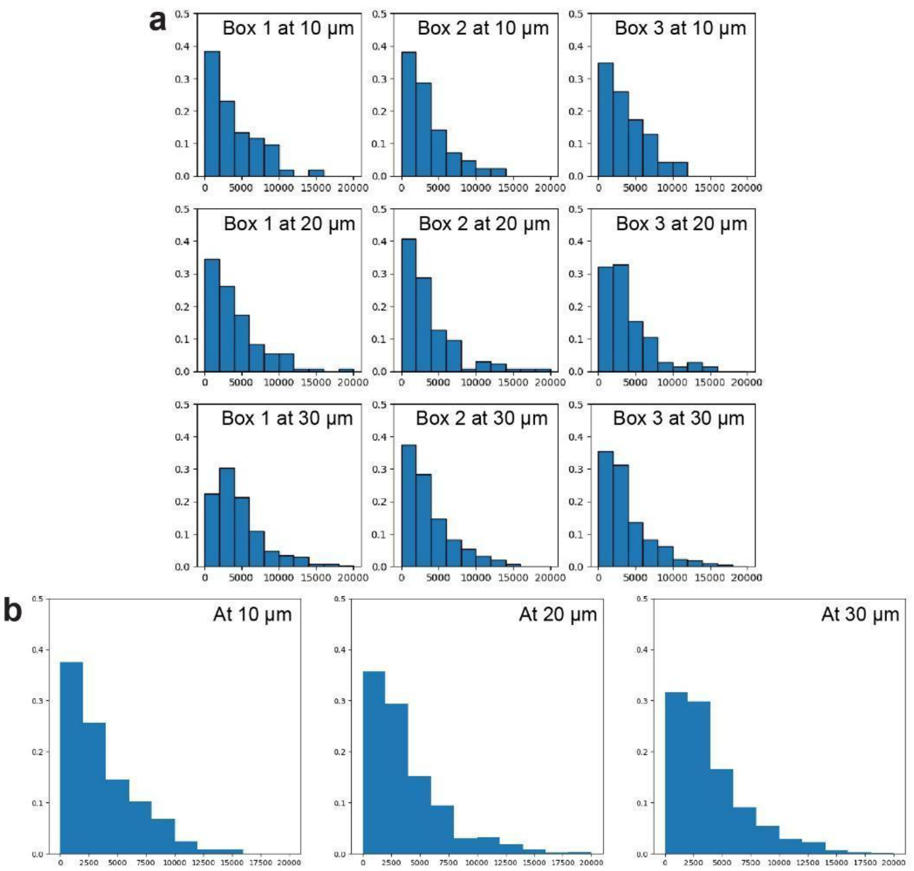


**Sup. Figure 10. Frequency distributions of the synapse volume (pre-synaptic + post-synaptic).**

**a**, Frequency distributions of synapses volumes (pre-synaptic + post-synaptic) of the nine boxes at 10 µm away, 20 µm away, and 30 µm away from the plaque. Synapse volume is shown in the unit of voxels. Each voxel equals 0.0000192 µm$^3$. **b**, Combined frequency distributions of synapse volumes (pre-synaptic + post-synaptic) at 10 µm away, 20 µm away, and 30 µm away from the plaque.

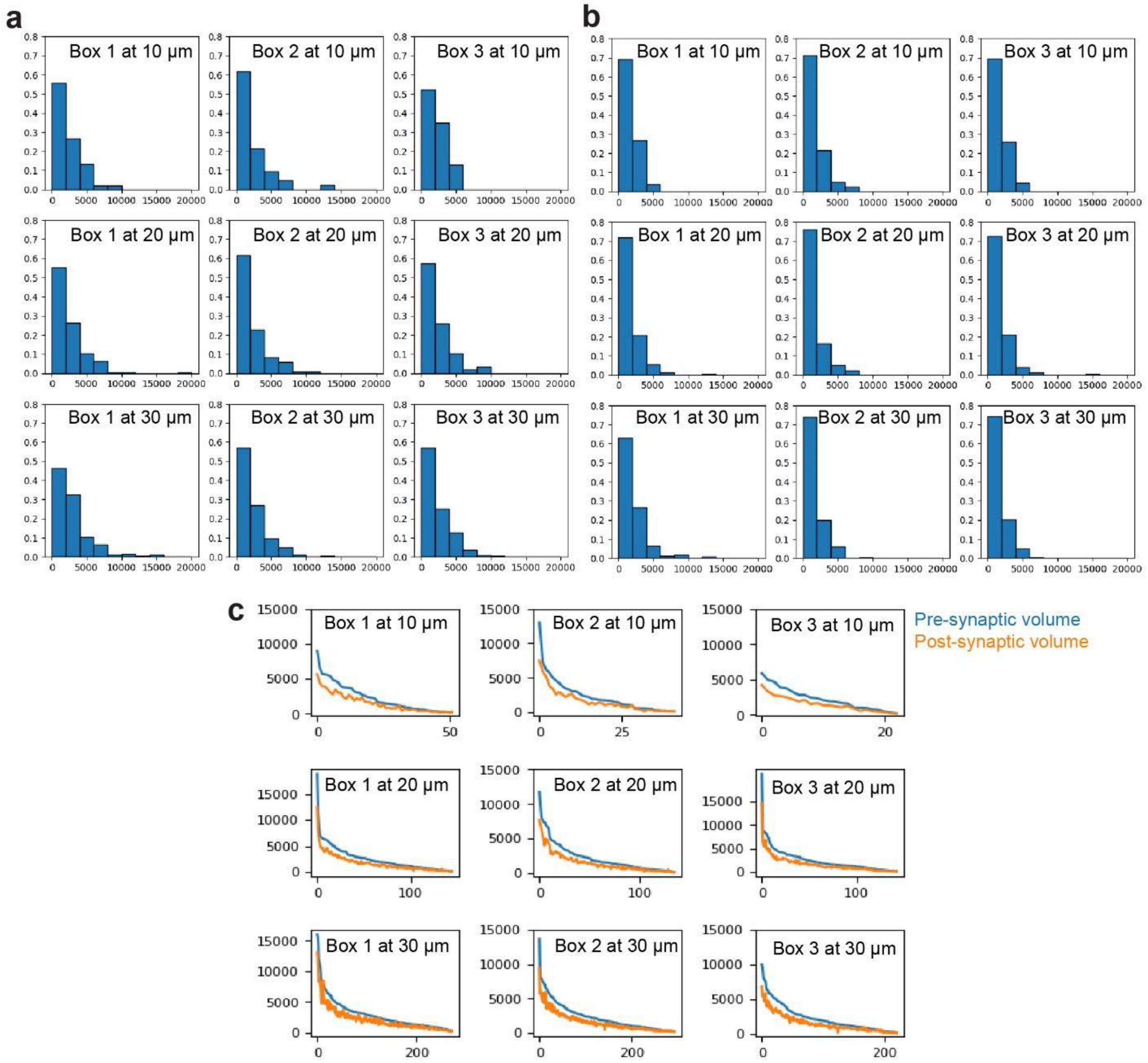


**Sup. Figure 11. Frequency distributions of the synapse volume (pre-synaptic or post-synaptic).**

**a**, Frequency distributions of synapse volumes (pre-synaptic) of the nine boxes at 10 μm away, 20 μm away, and 30 μm away from the plaque. Synapse volume is shown in the unit of voxels. Each voxel equals 0.0000192 $\mu m^3$. **b**, Frequency distributions of synapse volumes (post-synaptic) of the nine boxes at 10 μm away, 20 μm away, and 30 μm away from the plaque. Synapse volume is shown in the unit of voxels. Each voxel equals 0.0000192 $\mu m^3$. **c**, Comparison between pre-synaptic and post-synaptic volumes of every synapse in the nine boxes. x-axis, the number of synapses. y-axis, volume is shown in the unit of voxels. Each voxel equals 0.0000192 $\mu m^3$. Note the difference between the blue and orange curves.

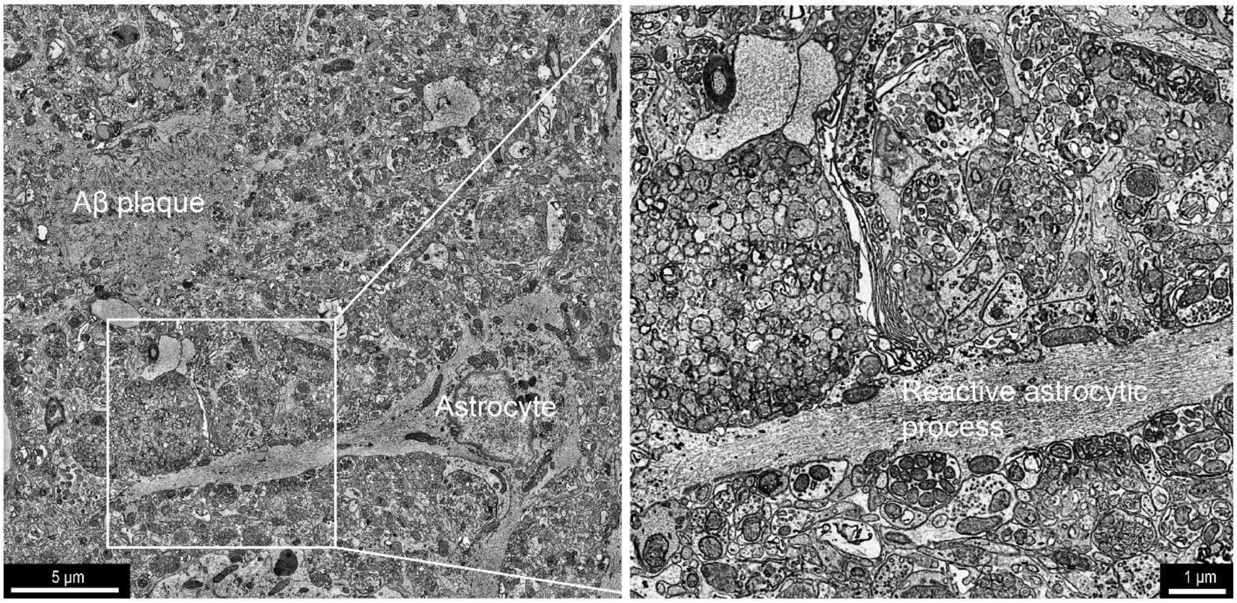


**Sup. Figure 11. An astrocyte and its reactive astrocytic process around an amyloid-β plaque.**

**Sup. Table 1. The final concentrations of nanobody probes used in the immunofluorescence of this work.**

| Target | Clone No. | Fluorescent dyes conjugated | Final conc. |
|---|---|---|---|
| **pTau** | A2 | 5-TAMRA, Alexa Fluor 594, Alexa Fluor 647 | 0.01 mg/ml |
| **Aβ 40/42** | R3VQ | 5-TAMRA, Alexa Fluor 647 | 0.01 mg/ml |
| **CD11b** | DC13 | Alexa Fluor 488 | 0.01 mg/ml |
| **GFAP** | E9 | 5-TAMRA | 0.01 mg/ml |
| **Ly-6C/6G** | 21 | Alexa Fluor 647 | 0.01 mg/ml |

**Sup. Table 2.The dilution ratios of primary antibodies used in the immunofluorescence used of this work.**

| Name | Fluorescence dye conjugated | Vendor | Catalog No. | Dilution Ratio |
|---|---|---|---|---|
| **Phospho-Tau (Ser202, Thr205) Monoclonal Antibody (AT8)** | n.a. | ThermoFisher | MN1020 | 1:100 |
| **Alexa Fluor 647 anti-β-Amyloid, 17-24 Antibody (4G8)** | Alexa Fluor 647 | BioLegend | 800718 | 1:50 |

**Sup. Table 3. The final concentrations of secondary antibodies used in the immunofluorescence used of this work.**

| Name | Fluorescence dye conjugated | Vendor | Catalog No. | Dilution Ratio |
|---|---|---|---|---|
| **Goat anti-Mouse IgG (H+L) Cross-Adsorbed Secondary Antibody** | Alexa Fluor 488 | ThermoFisher | A-11001 | 1:2000 |

### Supplementary File

### Nanobody Purification and Dye-Conjugation Protocol

#### Transfection

- Use the Expi293 expression system (ThermoFisher) and 293fectin transfection reagent (ThermoFisher, 12347019).
- Refer to the system-specific transfection protocol for detailed steps.
- For each transfection:
    - Use 100 x $10^6$ cells in 100 mL of Expi293 media.
    - Add 100 µg of plasmid DNA, which should be prepared using an endotoxin-free maxi-prep kit (Qiagen,12362 ).

#### Nanobody Purification

(For each 100 mL of Expi293 culture)

Buffers:

- Tris buffer: 50 mM Tris, 150 mM NaCl
- Elution buffer: 250 mM imidazole, 50 mM Tris, 150 mM NaCl
- Tris-glycerol buffer: 50 mM Tris, 150 mM NaCl, 15% glycerol (diluted from 50% glycerol)

Procedure:

1. Harvest the supernatant seven days after transfection:
    - Spin the culture media at 1500 rpm for 5 minutes. Repeat once to further clean the supernatant.
2. Prepare Ni-NTA agarose (Qiagen, 30210):
    - Shake well, then take 2 mL slurry and transfer it to a 15 mL tube.
    - Add Tris buffer to fill the tube and centrifuge at 600g.
    - Remove the supernatant by aspiration.
3. Resuspend the beads:
    - Add 2 mL Tris buffer to the tube (final volume: 4 mL).
    - Mix thoroughly and split 2 mL into each 50 mL tube containing ~50 mL of culture supernatant.
4. Incubate tubes on a rocker at 4°C for 1 hour.
5. Transfer the supernatant to a chromatography column (BIO-RAD, 7321010) and rinse the tube with additional Tris buffer.
    - Allow the buffer to flow by gravity.
6. When dripping stops, cap the column and add 750 µL of elution buffer.
    - Incubate for 5 minutes, then uncap and collect the effluent into an Amicon Ultra-15 Centrifugal Filter Unit (Millipore-Sigma, UFC901024).

7. Repeat the elution process twice more with 750 µL of buffer each time.
8. After the final elution, spin the filter unit at 4000 rpm for 20 minutes at 4°C.
    - Reduce the volume to ~0.5 mL.
9. Add Tris-glycerol buffer and repeat the centrifugation process twice more to reduce the volume to ~0.5 mL.
10. Agitate the liquid with a pipette to resuspend the protein and transfer it to a clean 1.5 mL tube.
11. Measure the protein concentration using a Nanodrop.
12. Store the purified nanobody at -20°C.

**Maleimide Dye-Peptide Conjugation**

1. Synthesize GGGC peptide (available from ThermoFisher, Genscript, or your institution's core facility).
2. Obtain maleimide dye (e.g. ThermoFisher or Lumiprobe).
3. Dissolve 10 mg GGGC peptide in 500 µL of 1x PBS.
4. Dissolve the maleimide dye in 50-100 µL of DMSO (depending on solubility).
5. Add the dissolved dye to the peptide solution, maintaining a 5–10:1 molar ratio of peptide to dye.
    - If dye precipitates, add more DMSO (keeping total DMSO concentration below 30%).
6. Mix thoroughly and shake at 800 rpm at 4°C overnight on a thermo mixer.
7. Use HPLC to purify the dye-peptide conjugate.
8. Freeze-dry the purified conjugate and re-dissolve it in dd$H_2O$ to a final concentration of 4 mM.
    - If insoluble, add up to 10% DMSO.
9. Store the conjugate at -20°C.

**Sortase Reaction**

1. Prepare a reaction mixture containing:
    - Nanobody, dye-peptide conjugate, sortase, and sortase buffer.
    - Refer to Fang, Tao, Xiaotang Lu, Daniel Berger, Christina Gmeiner, Julia Cho, Richard Schalek, Hidde Ploegh, and Jeff Lichtman. 2018. "Nanobody Immunostaining for Correlated Light and Electron Microscopy with Preservation of Ultrastructure." Nature Methods 15 (12): 1029–32. for protocol details.
    - A commercial sortase kit like the SiMPLe Protein Labeling Kit (BPS Bioscience, 79392) may also be used.
2. Incubate on a thermo shaker at 500 rpm and 12°C for 3 hours.
3. Prepare Ni-NTA agarose 15 minutes before the reaction ends:
    - Take 100 µL Ni-NTA, wash with Tris buffer (three times), and spin down at 600g.

- Leave a small amount of buffer after the final spin and transfer the beads to the reaction tube.

4. Cap and tape the reaction tube horizontally. Shake at 1000 rpm for 20 minutes.
5. Centrifuge briefly and transfer the supernatant to an Amicon Ultra-4 Centrifugal Filter Unit (Millipore-Sigma, UFC801024).
6. Rinse the reaction tube with 200 µL Tris-glycerol buffer and transfer to the filter unit.
7. Spin at 4000 rpm for 20 minutes at 4°C.
8. Add Tris-glycerol buffer and repeat the centrifugation twice, with a 30-minute spin on the final round.
9. Continue spinning until the volume is below 500 µL.
10. Transfer the final product to a 1.5 mL tube and measure the concentration using a Nanodrop at A280.
    - The final concentration should be 0.5–1 mg/mL.
11. Store the product at -20°C.
